# Large sharp-wave ripples promote hippocampo-cortical memory reactivation and consolidation

**DOI:** 10.1101/2025.06.27.662061

**Authors:** Heath L. Robinson, Ralitsa Todorova, Gergo A. Nagy, Anna Gruzdeva, Praveen Paudel, Azahara Oliva, Antonio Fernandez-Ruiz

## Abstract

During sleep, ensemble activity patterns encoding recent experiences are reactivated in the hippocampus and cortex. This reactivation is coordinated by hippocampal sharp-wave ripples (SWRs) and is believed to support the early stages of memory consolidation. However, only a minority of sleep SWRs are associated with memory reactivation in the hippocampus and its downstream areas. Whether that subset of SWRs have specific physiological characteristics and directly contribute to memory performance is not known. We identified a specific subset of large SWRs linked to memory reactivation in both the hippocampus and prefrontal cortex (PFC) of mice, and found that their occurrence selectively increased during sleep following new learning. Closed-loop optogenetic SWR boosting during sleep was sufficient to enhance ensemble memory reactivation in hippocampus and PFC. This manipulation also improved subsequent memory retrieval and hippocampal-PFC coordination during waking, causally linking both phenomena to SWR-associated ensemble reactivation during sleep.

## Introduction

New memories are consolidated during sleep following learning, a process that leads to the formation of stable, long-term representations^1,2^. The hippocampus plays a key role in the initial memory encoding ^3^. According to the influential systems memory consolidation theory, memories are gradually transferred from the hippocampus to the neocortex for permanent storage^4–6^. This process has been hypothesized to be mediated by synchronous network patterns generated in the hippocampus, known as sharp-wave ripples (SWRs)^7–10^. SWRs occur during non-rapid eye movement (NREM) sleep and waking immobility periods. They coordinate the reactivation of hippocampal neurons that encoded recent experiences and broadcast these memory representations to the neocortex and other brain areas^10–14^ . Numerous studies, including both correlational analyses as well as causal manipulations, have established the role of hippocampal SWRs in memory formation and consolidation^15–25^. In downstream structures such as the prefrontal cortex (PFC), SWRs entrain local circuits, driving the coordinated reactivation of experience-related activity patterns^26–30^. Furthermore, the reactivation of neural population activity in both the hippocampus and prefrontal cortex has been linked to novel experiences and memory demands^26,31–37^.

Despite this wealth of evidence, an apparent paradox is that only a small subset of sleep SWRs are linked to the reactivation of recent experiences. Most studies, using a variety of different methods, converge to only ∼10 to 30 % of SWRs significantly associated with significant memory reactivation or “replay”^12,15,20,25,34,36,38^. This fraction could be even smaller for memory reactivation in downstream structures involved in memory consolidation, such as the PFC. These observations raise the question of whether the subset of SWRs that successfully elicit memory reactivation in the neocortex, and thus presumably contribute to memory consolidation, have unique physiological characteristics or whether their impact is determined by some additional variables that converge with SWRs. In this study, we aimed to distinguish between these two hypotheses and establish a direct link between SWR features and memory performance. Furthermore, studies in humans have shown that increased functional coordination between the hippocampus and cortex during rest following new learning correlates with successful memory recall ^39,40^. However, whether sleep SWRs mediate a persistent enhancement of hippocampo-cortical coordination and subsequent memory recall is not known.

To answer these questions, we performed neural ensemble recordings in the hippocampus and PFC of mice during a memory task and sleep, together with closed-loop optogenetic manipulation of SWRs. We found that only a specific subset of large SWRs were associated with memory reactivation in both structures. The rate of these large SWRs selectively increased in sleep after learning and correlated with successful memory performance. To directly test their contribution to memory, we optogenetically boosted sleep SWRs following a learning paradigm otherwise insufficient for mice to consolidate a new experience. SWR boosting enhanced memory reactivation in both the hippocampus and PFC and, remarkably, allowed mice to successfully recall the new experience after sleep. Oscillatory synchrony between hippocampus and PFC during memory recall was also enhanced, linking SWR-mediated sleep memory reactivation to persistent functional connectivity changes, in agreement with similar reports from human memory studies.

## Results

### Large SWRs are associated with stronger memory reactivation in CA1 and PFC and modulated by learning

We implanted mice with high-density silicon probes in the dorsal hippocampus (CA1 region) and PFC (pre-limbic region) (Fig. 1A and Fig. S1). We simultaneously recorded large ensembles of cells in both regions (n = 1827 CA1 and 1103 PFC neurons from 21 mice) while mice performed a hippocampal-dependent memory task (object displacement, see Methods) and in sleep before and after the task. To analyze memory reactivation, we employed an unsupervised method (based on principal component analysis -PCA- followed by Independent Component Analysis -ICA-) to detect the synchronous coactivity of groups of neurons, or cell assemblies, during learning, as in previous work ^23,24,41,42^. We then compared whether these task-related cell assemblies in PFC and CA1 reactivated during SWRs in subsequent sleep. Using this approach, we found that 35.4% of SWRs were associated with significant memory reactivation in the hippocampus and 19.7% in PFC (P < 0.05, n = 353.600 SWRs from 21 mice). The probability of PFC reactivation was higher for SWRs associated with significant CA1 reactivation (Fig. S2A).

**Figure 1:**
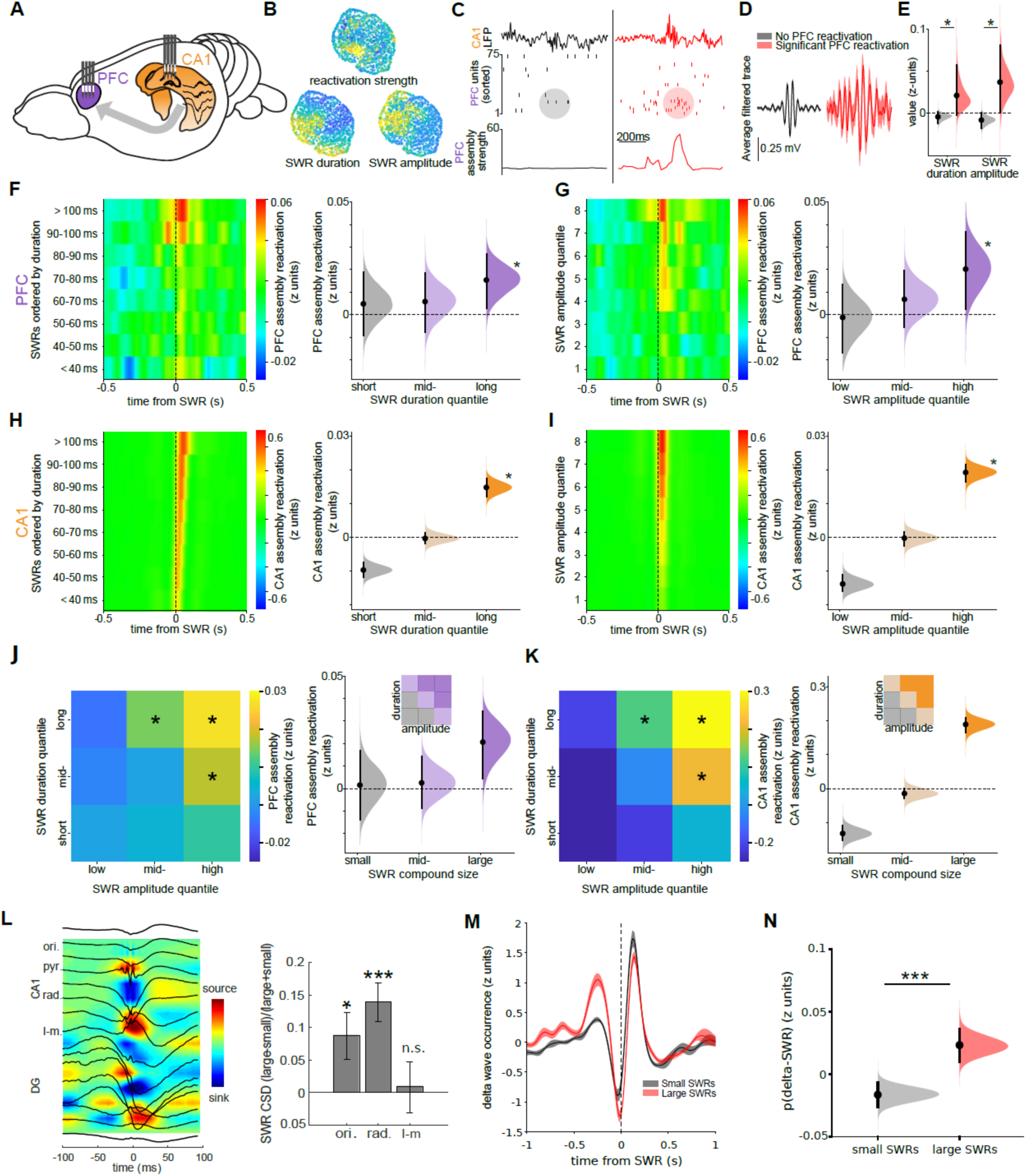
Enhanced memory reactivation in hippocampus and cortex during a subset of SWRs **A)** Schematic of recording design. **B)** Visualization of SWR features using a 3D UMAP embedding. Each point is a single SWR from an example session, color coded according to different features. **C)** Examples of SWR without (left) and with (right) associated PFC cell assembly reactivation. Top: CA1 LFP traces. Middle: raster plots of neuronal firing, a colored circle indicates assembly activity accompanying the hippocampal SWR; Bottom: assembly reactivation strength. **D)** Average filtered CA1 LFP trace for SWRs associated with (red trace) or without (black trace) significant reactivation of PFC cell assemblies. Data is from the same session as B-C. **E)** SWRs associated with significant reactivation of PFC cell assemblies were longer (P = 0.043, n = 69,522 SWRs, hierarchical bootstrap test) and had higher amplitude (P = 0.025, n = 284,078) than the remaining SWRs. Black dots: mean; black lines: 95% confidence intervals; curves: the hierarchically resampled mean distributions. **F)** Left: reactivation strength of PFC assemblies around SWRs sorted by duration. Right: hierarchically bootstrapped estimates of the PFC assembly reactivation during SWRs classified by duration quantiles (P = 0.25 / 0.20 / 0.0104 for short, mid-length and long ripples respectively, hierarchical bootstrap test). **G)** Same as F but with SWRs sorted by amplitude (P= 0.55 / 0.14 / 0.013 for low, mid and high-amplitude SWRs respectively). **H)** Same as F but for CA1 assemblies (P=1 / 0.65 / 0 for short, mid-length and long SWRs respectively). **I)** Same as H but with SWRs sorted by SWR amplitude (P=1 / 0.57 / 0 for low, mid and high-amplitude SWRs). **J)** Left: PFC assembly responses to SWR quantile grid split according to both duration (y-axis) and amplitude (x-axis). Note that PFC responses were significant only for SWR quantiles of long-duration (top) and high-amplitude (right) (*P<0.05, Wilcoxon signed-rank tests). Right: Hierarchically bootstrapped estimates of PFC reactivation strength. Note that small and mid-length SWRs showed no significant PFC reactivation (P = 0.42 / 0.34, hierarchical bootstrap test), but large SWRs were followed by significant reactivation (P = 0.0076). **K)** Same as J but for CA1 assemblies (P = 1 / 0.96 / 0 hierarchical bootstrap test for small, mid-size, and large SWRs respectively). **L)** Left: Averaged CSD profile of SWRs during a representative session. Silicon probe shank spans CA1 and DG layers (ori = stratum orients, rad. = str.radiatum, l-m. = str. lacunosum-moleculare) Right: quantification of relative current sink magnitude per layer for large versus small SWRs (*/*** P < 0.05/ 0.001, signed-rank test; n = 15 sessions from 5 mice).**M)** Mean peri-event time histogram of delta wave occurrence (top) and spindle occurrence (bottom) around small (black) and large SWRs (red). Mean ± SEM. **N)** Probability of a delta wave occurring before SWR (-400 to -100 ms) was higher for large SWRs than small SWRs (P = 0, hierarchical bootstrap test; N=18 sessions in 10 animals)

We asked whether SWRs that elicited ensemble reactivation in downstream areas, such as the PFC, and could therefore promote memory consolidation, have specific defining features. To gain intuition about which features may be more relevant, we first represented all SWRs as points in a high- dimensional space constructed using different features (see Methods) and embedded them in a low- dimensional space using Uniform Manifold Approximation and Projection (UMAP) for visualization. This method revealed that SWR properties such as amplitude, duration or frequency varied along a continuum (Fig. 1B and Fig. S2) ^36,43^ . Color-coding SWRs based on the reactivation strength of concurrent PFC ensemble activity suggested a correlation between SWR amplitude and duration with PFC reactivation strength (Fig. 1B). Classifying all sleep SWRs in two binary categories, those with and without significant associated memory reactivation in PFC, confirmed that the former had significantly larger duration and amplitude (Fig.1C-E), but did not significantly differ in other features (Fig. S2).

To examine in more detail the relationship between memory reactivation strength and SWR duration and amplitude, we classified SWRs based on these features and calculated peri-SWR histograms of PFC assembly reactivation strength for each of the categories. Only the topmost duration and amplitude SWR quantiles were associated with strong PFC memory reactivation (Fig. 1F-G). We obtained similar results when we performed the same analysis with CA1 assembly reactivation strength (Fig. 1H-I). However, in the case of CA1, reactivation strength increased more gradually with increasing SWR duration and amplitude in contrast to a more abrupt increase in PFC.

SWR duration and amplitude were only moderately correlated (Fig. S2), and our initial observations suggested that only a subset of SWRs with both large amplitude and duration were associated with strong memory reactivation (Fig. 1B). Therefore, we classified SWR into groups of increasing joint amplitude and duration. Memory reactivation strength in both the PFC (Fig. 1J) and CA1 (Fig. 1K) was largest for SWRs in the top duration and amplitude quantile, but weak for SWRs with large amplitude but short duration or long duration but low amplitude.

We further examined whether small and large SWRs differ in their relationship with upstream or downstream regions. First, we analyzed the relative strength of synaptic inputs during small and large SWRs by performing current source density analysis (CSD) on hippocampal laminar recordings. Both groups of SWRs had qualitatively similar CSD profiles, suggesting that they were generated by the same synaptic inputs. However, large SWRs were associated with stronger stratum oriens and, specially, stratum radiatum current sinks compared to small SWRs (Fig. 1L). In contrast, there were no significant differences in the magnitude of stratum lacunoso-moleculare sinks (Fig. 1L). This result suggests that an enhanced CA3 input (that primarily targets stratum radiatum) may be the main driver of large SWR generation.

Next, we analyzed the relationship between hippocampal SWR and cortical sleep oscillations. We focused on delta waves and spindles, which coupling with SWRs supports memory consolidation^44,45^. Small and large SWRs alike tend to be followed by PFC delta waves, but large SWRs were significantly more likely than small SWRs to follow delta waves (Fig. 1M-N and Fig. S3), that is to occur at the onset of the cortical UP state. In addition, large SWRs were also more likely to be coupled with PFC spindles than small SWRs (Fig. S5).

Finding that memory reactivation was associated with a specific subset of larger SWRs prompted us to ask whether the occurrence of these SWRs was correlated with successful memory consolidation. We tested mice in two versions of the object displacement task (Fig. 2A) ^46^ . In the first one, mice were allowed to explore two identical objects for three 5-minute trials. For the memory recall test, conducted after a 4-hour sleep session in their home cages, one object was displaced to a different location. Mice displayed an increased exploration of the displaced object as quantified by a discrimination index (Fig. 2B and Fig. S4), establishing that they correctly remembered its original location. However, when mice were only allowed to explore the object for one 5-minute trial, they did not show an exploratory bias for the displaced object during the following test (Fig. 2B and Fig. S4). Mice did show an increased exploration of the displaced object after 5-minute exploration when the test was conducted only 1 hour later (Fig. S4C). This suggests that the reduced experience during 5-minute sessions was enough to support short-term memory recall but not enough for longer-term memory consolidation. Sleep structure and the amount of NREM were similar in the rest periods after both versions of the task (Fig. S5).

**Figure 2:**
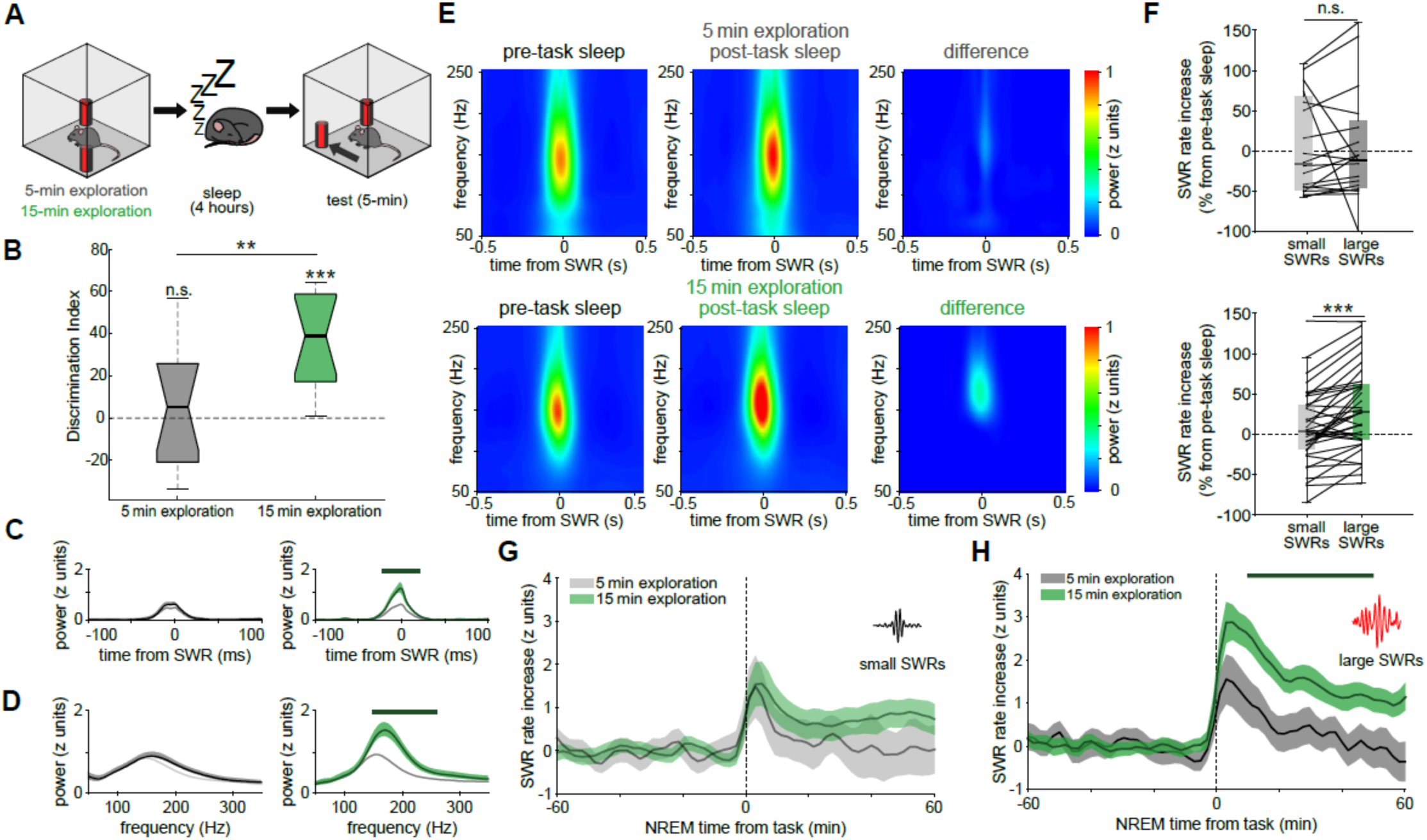
Increased fraction of large SWRs correlates with memory performance. **A)** Animals were trained on the object displacement memory task, with either a single or three 5- minute exploration sessions. **B)** Memory performance (object discrimination index) on the test session was significant for the 15-minute (n = 13 mice, P = 2.4e-4, Wilcoxon signed rank test) but not the 5- minute group (n = 13 mice, P = 0.68, Wilcoxon signed rank test; P = 0.011 for 5 vs 15-minute group, Wilcoxon rank-sum test). **C)** Spectral power within the ripple band (150-250 Hz) increased for SWRs following 15-minute but not 5-minute exploration (n = 38 and 20 sessions, respectively). A line indicates the significant interval (P < 0.05, Monte Carlo test). **D)** Spectral profile of SWRs (within 50 ms of the SWR peak) was unaffected after 5-minute exploration (left), while power in the ripple frequency band was enhanced following 15-minute exploration (P < 0.05, Monte Carlo test, n = 20 / 38 sessions); a line indicates the significant interval **E)** Average sleep SWR wavelet spectrograms for pre (left), post-task sleep (middle) and their difference (right), following either 5-minute (top, n = 20) or 15-minute (bottom, n = 38) exploration sessions. **F)** Fraction of large SWRs increased following 15-minute (bottom), but not 5-minute (top), object exploration sessions (5-minute: P = 0.68, n = 20 and 15- minute: P = 8.9×10^-6^, n = 38, Wilcoxon signed-rank test). **G)** Incidence of small SWRs in NREM (one hour of contiguous NREM sleep before and after the task) was similar between 5-minute and 15- minute training conditions (P > 0.05, hierarchical bootstrap test). **H)** Incidence of large SWRs in post-task NREM sleep increased following 15-minute exploration (P < 0.05); a line indicates the significant interval. C-G show mean ± SEM.

We next compared SWR features in the sleep sessions before and after the initial object exploration. SWRs before and after the 5-minute exploration sessions were similar in amplitude and duration (Fig. 2C-E). In contrast, SWRs following 15-minute exploration sessions were larger on average than in the preceding sleep period (Fig. 2C-E and Fig. S6). We thus divided all SWRs into “large” and “small” using a compound measure combining duration and amplitude (see Methods) and calculated their relative fraction in each sleep period. Overall SWR rate (0.43 and 0.47 Hz in pre and post sleep respectively, p = 0.079, Wilcoxon signed-rank test) and the relative fraction of small and large SWRs did not change following 5-minute object exploration sessions (Fig. 2F). Following 15-minute exploration sessions, the overall SWR rate increased (0.46 and 0.52 Hz, p = 6.2e-4, Wilcoxon signed- rank test), specifically due to a larger fraction of large, but not small, SWRs (Fig. 2F). Furthermore, there was a significant positive correlation between the post-sleep fraction of large SWRs and task recall performance after 15-minute but not 5-minute exploration sessions (Fig. S7).

We then examined the temporal dynamics of SWR occurrence. SWR rate and fraction of large SWRs after 15-minute exploration increased sharply at the start of the post-learning NREM sleep and then gradually decayed but remained high for ∼1 hour of NREM (Fig. 2G-H). Following the 5-minute exploration, only a transient increase of SWR rate was observed right at the start of NREM sleep, and no specific increase of large SWR occurrence (Fig. 2G-H). Moreover, SWR rate already increased gradually during immobility task periods during 15-minute training sessions compared to 5-minute training sessions (Fig. S4D-E).

### Closed-loop optogenetic SWR boosting enhanced memory reactivation and recall performance

After describing a correlation between large SWRs during sleep and successful memory retrieval, we next wanted to establish whether there is a causal link between large SWRs and memory performance by specifically manipulating them during the post-task sleep period. For this purpose, we expressed channelrhodopsin (ChR2) in CA1 pyramidal cells using viral and transgenic approaches (see Methods). Mice were implanted with silicon probes (or wire arrays) in the hippocampus and the prefrontal cortex and bilateral optic fibers above CA1 (Fig. 3A). We detected SWRs in real-time during post-task sleep sessions and optogenetically boosted them using a closed-loop approach (Fig. 3B and Fig. S8). A 100 ms low-intensity blue-light pulse delivered upon SWR detection, prolonged the ongoing oscillation and associated spiking activity (Fig. 3C). This manipulation also resulted in a boost of SWR power (Fig. 3D). To test whether this manipulation could enhance memory performance, we applied it during the sleep period following 5-minute object exploration sessions. SWR boosting resulted in an increase in the total fraction of large SWRs during post-task sleep similar to that naturally occurring following 15-minute exploration sessions (Fig. 3E). Remarkably, while after 5-minute exploration sessions without optogenetic stimulation mice did not recall the displaced object, memory recall was successful in sessions with SWR boosting during sleep (Fig. 3F). This result indicates that optogenetic boosting of sleep SWR was able to enhance memory performance.

**Figure 3:**
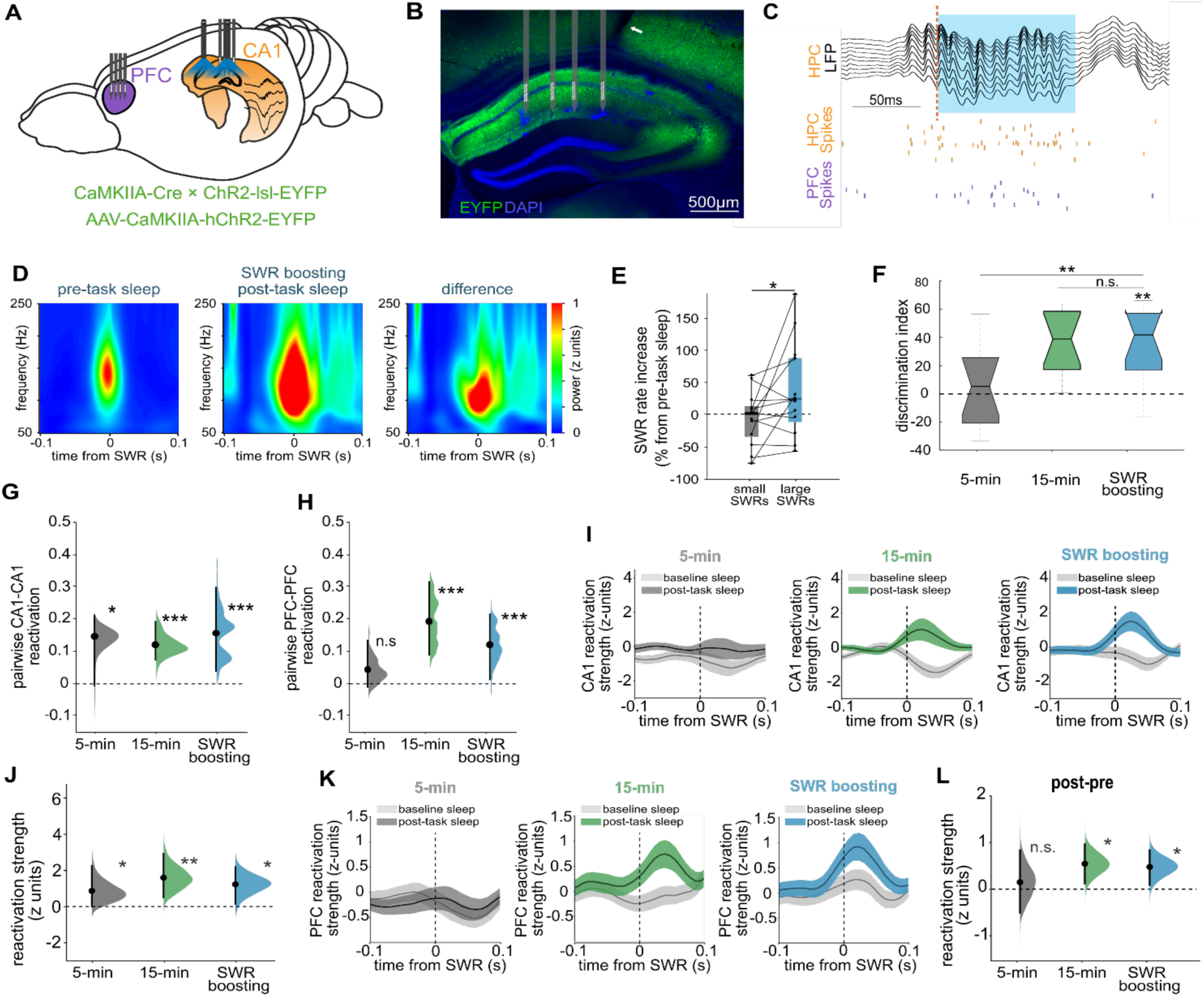
Optogenetic SWR boosting enhances memory reactivation and performance. **A)** Schematic of CA1 and PFC recordings combined with optic fibers targeting the hippocampus bilaterally. **B)** Histological section showing the expression of ChR2-EYFP (green) in pyramidal cells; white arrow: optic fiber track. **C)** Example of optogenetic SWR boosting. Black trace is CA1 LFP, vertical dashed bar shows SWR real-time detection and blue rectangle shows optogenetic stimulation pulse. Raster plot below shows CA1 (in gold) and PFC (in purple) units firing. **D)** Average sleep SWR wavelet spectrograms for pre (left) and post-task sleep (middle) with SWR boosting (n = 13 mice; 120 -230 Hz spectral power comparison between post-task and baseline sleep, P < 0.05, Monte Carlo test). **E)** SWR boosting resulted in an increased incidence of large SWRs in post-task sleep (*P = 0.028, n = 13, Wilcoxon signed-rank test). **F)** Memory recall performance (object discrimination index) for no stimulation 5-minute and 15-minute training sessions (same data as Figure 2B) and SWR boosting sessions (n = 13 mice, P = 1.7e-3, Wilcoxon signed rank test. SWR boost vs 5-minute P = 8.9e-3, Wilcoxon rank sum test. SWR boost vs 15-minute P = 0.96). **G)** CA1 cell pair co-firing in SWRs (pairwise reactivation). Grey: 5-minute exploration (P = 0.039, n = 18,708 cell pairs, hierarchical bootstrap test); green: 15-minute exploration (P = 0, n = 16,215); blue: 5-minute exploration followed by SWR-boosting (P = 0.001, n = 10,963); black dots: mean estimates; black lines: 95% confidence intervals; curves: the resampled mean distributions. **H)** Same as G) but for PFC. 5-minute exploration (grey, P = 0.14, n = 12,739 pairs); 15-minute (green: P = 0, n = 10,729); SWR-boosting (blue: P = 0.020, n = 11,157). **I)** Averaged peri-SWR CA1 assembly reactivation (using PCA-ICA) strength following 5-minute exploration (left), 15-minute (middle), and 5-minute followed by SWR-boosting (right); light grey: baseline pre-task assembly activity. Mean ± SEM. **J)** Quantification of I) measured by hierarchical bootstrap estimate of assembly reactivation in post-task sleep relative to baseline sleep: 5-minute exploration (grey: P = 0.032, n = 97 assemblies); 15-minute (green: P = 1.8e-3, n = 189); SWR boosting (blue: P = 0.016, n = 131 assemblies). **K-L)** Same as I-J) but for PFC assemblies. 5-minute (grey: P = 0.35, n = 79 assemblies); 15-minute (green: P = 0.0104, n = 96), SWR-boosting (blue: P = 0.011, n = 115).

To verify whether the behavioral impact of SWR boosting depends on the temporal specificity of the manipulation we performed a control experiment. We delivered the same optogenetic stimulation but after a random delay upon SWR detection in post-task sleep following 5-minute training sessions. This manipulation elicited a similar physiological response (evoked ripple-like oscillations and associated spiking in CA1) than the SWR timed stimulation (Fig. S8). In contrast, delayed stimulation did not have any significant effect on memory performance (Fig. S8), suggesting that the enhancement previously observed relies on precisely timed stimulation of CA1.

We hypothesized that the observed differences in memory performance following 5 or 15-minute exploration and the behavioral effects of optogenetic SWR boosting could be explained by differences in post-task neural reactivation in CA1 and PFC. To test that, we assessed sleep memory reactivation using several approaches. First, we measured the correlation of neural firing patterns in pairs of cells between the task and post-sleep SWRs. Pairwise reactivation was significant (relative to baseline sleep) in CA1, but not in the PFC following 5-minute exploration (Fig. 3G-H). In contrast, after both 15-minute or 5-minute exploration followed by SWR boosting, we observed significant memory reactivation in both CA1 and PFC (Fig. 3G-H).

Next, we examined the reactivation of CA1 and PFC cell assemblies in sleep SWRs. Cell assemblies in CA1 only showed a weak reactivation following 5-minute exploration, and PFC assemblies displayed no significant reactivation at all (Fig. 3I-L). In contrast, we observed robust significant assembly reactivation in both structures after 15-minute exploration sessions, as well as 5-minute exploration with SWR boosting (Fig. 3I-L). To control for potential confounding effects due to differences in firing rate, we repeated these analyses after subsampling to match within-SWR firing rates in pre and post-task sleep for all conditions. The analysis resulted in similar enhanced assembly reactivation after 15-minute training and with SWR boosting (Fig. S9). Furthermore, we observed a session-wise positive correlation between the rate of optogenetically boosted SWRs and memory performance (Fig. S7). Optogenetic stimulation did not alter overall sleep structure or the relative fraction of NREM sleep epochs (Fig. S5). However, SWR boosting did enhance the coupling of hippocampal SWR and PFC delta waves, in a similar manner to the effects of extended training (Fig. S3). Overall, these results suggest that the improvement in memory performance due to optogenetic SWR boosting was due to an enhancement of task-related neural reactivation in CA1 and PFC.

### Enhanced CA1-PFC coordination during memory consolidation and recall

Recent work in humans has shown that functional connectivity between the hippocampus and cortex increases after learning, and that this increase correlates with subsequent memory recall performance ^39,40^. We thus hypothesized that the coordination between CA1 and PFC during sleep would increase following 15-minute but not 5-minute exploration sessions, and that a similar effect would be induced by optogenetic SWR boosting. We performed two different analyses to test this hypothesis.

First, we examined the coordinated reactivation of CA1 and PFC cell assemblies. PFC cell assemblies increased their reactivation rate in SWRs associated with significant CA1 reactivation in sleep following 15-minute exploration, and after 5-minute exploration followed by sleep SWR boosting, but not after 5-minute exploration without stimulation (Fig. S10A). Moreover, we found evidence for cross- regional assembly–assembly-specific coordination as PFC assemblies consistently reactivated with specific CA1 assemblies throughout sleep sessions following 15-minute training or 5-minute exploration followed by SWR boosting (Fig. S10).

To test for CA1-PFC coordination directly, we investigated the coordinated reactivation of pairs of CA1 and PFC cells in SWR before and after the task. CA1-PFC pairwise reactivation increased following 15-minute but not 5-minute exploration sessions (Fig. 4A). SWR boosting after 5-minute exploration also resulted in enhanced pairwise cross-regional reactivation (Fig. 4A). Next, we identified cross- regional assemblies, synchronous groups of coactive cells with members from both CA1 and PFC (Fig. 4B)^29,47^, and assessed their reactivation in sleep SWR before and after the task. CA1-PFC cell assemblies showed significant coordinated reactivation after 15-minute exploration or 5-minute exploration followed by sleep SWR boosting, but not after 5-minute exploration without stimulation (Fig. 4C).

**Figure 4:**
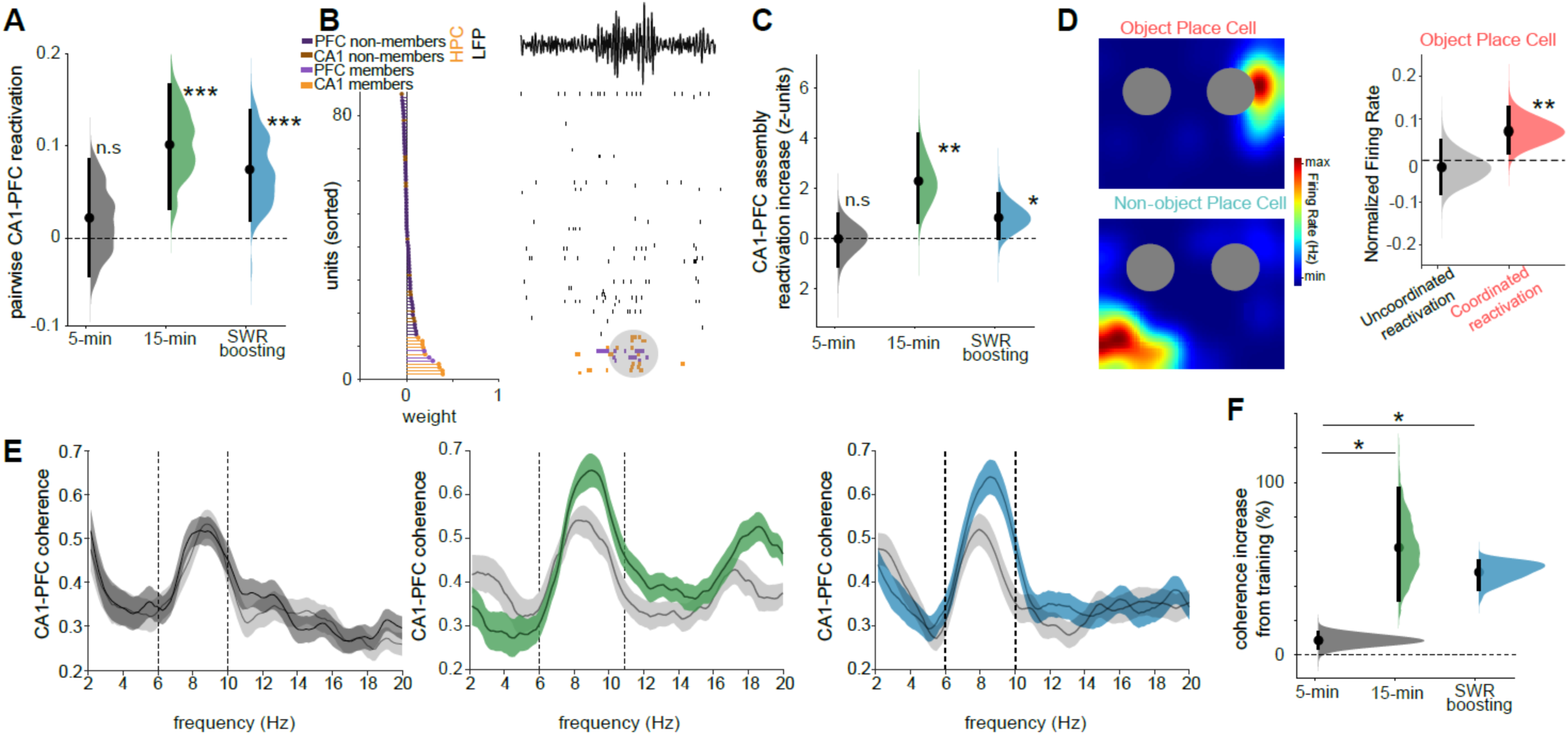
Coordination of CA1 and PFC activity during sleep and memory recall **A)** Hierarchically bootstrapped estimates of the CA1-PFC cell pair cofiring (pairwise reactivation): 5- minute exploration (grey: P = 0.32, n = 10,342 cell pairs; hierarchical bootstrap test); 15-minute (green: P = 0, n = 4,573); 5-minute exploration followed by SWR-boosting (blue: P = 0.012, n = 16,334 cell pairs). **B)** Example CA1-PFC assembly. Left: CA1 and PFC units sorted by their contribution to selected assembly (assembly members in bright color at the bottom). Right: examples of raster plots of unit firing showing assembly reactivation (shaded ovals) during a SWR (CA1 filtered LFP on top). **C)** Quantification of peri-SWR CA1-PFC assembly reactivation strength (using PCA-ICA, see Methods) measured by a hierarchical bootstrap estimate of assembly reactivation in post-task sleep relative to baseline sleep: 5-minute (grey, P = 0.032, n = 29 assemblies); 15-minute (green, P = 1.8e- 3, n = 28 assemblies); 5-minute with SWR boosting (blue, P = 0.016, n = 26 assemblies). **D)** Left: Rate map examples of an Object Place Cell (top) and a Non-object Place Cell (bottom). Objects to scale as gray circles. Right: Object Place Cells fired more during CA1-PFC coordinated reactivation events than during uncoordinated reactivation ones in post-task sleep (P = 0.0052, Wilcoxon rank-sum test). **E)** CA1-PFC LFP coherence in training and test. Left: CA1-PFC coherence was similar during 5-minute exploration (light grey) and the subsequent test session (black). 15-minute exploration (middle) and SWR boosting following 5-minute exploration (right) resulted in enhanced CA1-PFC coherence in the subsequent test session (green or blue) compared to training (light grey). Mean ± SEM. Dashed vertical lines indicate theta frequency band.**F)** Quantification of D) measured by a hierarchical bootstrap estimate of the increase in hippocampo-prefrontal coherence during test, normalized by coherence during training (5-minute versus 15-minute training, P = 0, n = 9 and 13 sessions respectively; 5-minute versus 5-minute training with SWR boosting, P = 0, n = 9 / 5 sessions; 15-minute versus 5-minute training with SWR boosting P = 0.24, n = 13 / 5 sessions).

Given the observed increase in coordinated CA1-PFC reactivation following extended experience as well as SWR boosting, we examined the representational content of coordinated cross-regional reactivation events (SWRs with significant reactivation of both CA1 and PFC assemblies). We classified all hippocampal spatially selective cells into those that encoded object locations (‘object place cells’) or other locations in the maze (‘non-object place cells’) (Fig. 4D). Object, but not non- object, place cells selectively increased their firing rate in SWRs associated with significant coordinated CA1-PFC reactivation (Fig. 4D).

Finally, we asked whether the enhanced cross-regional coordination during sleep persisted during waking memory recall. To answer this, we measured CA1-PFC theta frequency LFP coherence, as it has been previously shown to be associated with cross-regional communication and successful memory performance^48,49^. We found that CA1-PFC theta coherence during the test phase was largely increased compared to the initial object exploration in both 15-minute sessions and 5-minute sessions followed by SWR boosting, in contrast to 5-minute exploration sessions (Fig. 4E-F). This observation suggests that extended training and artificial SWR boosting result in an enhanced CA1-PFC coordination that supports successful memory recall.

## Discussion

Neural ensemble reactivation in the hippocampus and associated cortical areas, such as the PFC, is regarded as a cellular hallmark of memory consolidation. This process is primarily associated with a minority of SWRs (∼10-20%) during NREM sleep^11–13,15,33^. SWRs are highly heterogeneous network events, as characterized in previous studies^21,50–52^. However, the significance of SWR heterogeneity in memory consolidation has not been established. We found that the subset of SWRs that were associated with the reactivation of a hippocampus-dependent memory task in CA1 and PFC were those with both the largest duration and amplitude. The fraction of these large SWRs selectively increased during sleep following new learning, but only when learning experience was sufficient for successful memory retrieval 4 hours later. Previous work reported global increases in SWR rate during sleep following learning^23,46,53^ and an association of SWR duration and amplitude with increased memory load during learning^21,54,55^. Our results expand these earlier observations and demonstrate that large SWRs promote ensemble reactivation in the hippocampus and downstream areas in support of memory consolidation.

To move beyond correlational observations and establish a more direct link between SWRs, ensemble reactivation and memory performance, we deployed a closed-loop optogenetic stimulation approach^21^. Brief optogenetic activation of CA1 pyramidal cells triggered by real-time detection of sleep SWRs boosted them, mimicking the effect of extended learning on SWR features. Optogenetically boosted SWRs resulted in enhanced memory reactivation in both CA1 and PFC, demonstrating a direct link between SWR features and the associated dynamics of neuronal population spiking activity. Remarkably, this manipulation, but not the same stimulation delivered after a random delay upon SWR detection, improved mice memory performance, enabling them to successfully recall a brief prior experience, normally insufficient for successful retrieval. This effect was similar to that of extended learning experience, suggesting that sleep memory reactivation can compensate for reduced learning time or promote the consolidation of weakly encoded experiences, in line with theoretical proposals on the role of “offline” memory replay for training artificial and biological neural networks^56–58^. During extended experience the rate of SWRs gradually increased. We hypothesized, in agreement with previous findings^32,59^, that awake reactivation can ‘tag’ specific neural ensembles and prioritize their subsequent reactivation during sleep, leading to enhanced consolidation. This effect could be preferentially driven by large SWRs, as we found they promote enhanced memory reactivation within and beyond the hippocampus. In addition, the specific sub-state within sleep when SWR occurs can also influence their role in memory processes, as suggested by a previous study^36^.

In a previous study^21^, we used the same closed-loop optogenetic approach to boost SWRs during the performance of a spatial short-term memory task, resulting in an improvement of learning performance.The present study extend those results indicating that large SWRs during sleep have an instrumental role in memory consolidation and, crucially, demonstrating that properly timed optogenetic stimulation can enhance neural ensemble reactivation in hippocampus and downstream structures. An important feature is that our optogenetic stimulation was delivered ∼20 ms after the start of the SWR. Due to the strong recurrent dynamics of the hippocampal circuit, once a given neural pattern has started to reactivate, increasing the excitation in the network (even if by artificial means) may boost the ongoing reactivation (e.g., by further depolarizing neurons that were getting subthreshold inputs), rather than recruit a different neural ensemble. This feature may also make it challenging to impose an arbitrary artificial pattern in the network, e.g. stimulating specific neurons to induce plasticity and generate a specific coactivity or sequential activity motif.

Multiple brain-wide phenomena, such as cortical oscillations and brain-state fluctuations, correlate with SWRs and are associated with memory consolidation^36,44,45,60–63^. Our results suggest that the contribution of some of these processes to memory could be mediated by their modulation of SWR features that, in turn, have a direct effect in promoting neural ensemble reactivation. For example, changes in neuromodulatory tone such as a decrease in the level of acetylcholine release in the hippocampus could create permissive substates when large SWRs are preferentially generated and memory reactivation is promoted.

A leading mechanism for the systems consolidation of new memories is the coordination of hippocampal and cortical activity promoted by SWRs^8,9,45,62,64,65^. Most of the evidence supporting this proposal is based on the coupling of hippocampal and cortical network patterns or mesoscopic activity^9,44,45^. Along the same lines, we found that large SWRs are more likely to follow PFC delta waves, suggesting that the onset of cortical UP states may be a privileged window for cortico-hippocampal interactions and influence memory consolidation. In comparison, much less is known about how the millisecond timescale dynamics of cell assemblies across structures changes during consolidation and how sleep processes are related to persistent cross-regional coordination during waking. We found that CA1-PFC spiking coactivity patterns increased their coordinated reactivation following extended learning experience, and correlated with successful memory recall. Furthermore, coordinated CA1-PFC reactivation events were more likely to encode object locations than other maze areas, suggesting that they play an important role in the consolidation of behavioral relevant memories.

During waking memory recall, the PFC and hippocampus exhibit enhanced coherence in the theta frequency band^48,49,66^. We found similar enhanced theta coherence following extended learning experience. Importantly, optogenetically boosting SWRs in sleep following reduced learning experience produced similar effects to extended training, that is increased CA1-PFC cell assembly reactivation during sleep and enhanced theta coherence during subsequent awake recall. This result provides a missing link between the functional coordination of hippocampus and cortex during sleep SWRs and persistent changes in cross-regional synchronization during waking. This finding offers a potential mechanism to explain observations in humans that found enhanced hippocampo-cortical functional connectivity following learning and its correlation with memory performance.

## Resource availability

### Lead Contact

Further information and requests for resources and reagents should be directed to and will be fulfilled by the Lead Contact, Dr. Antonio Fernandez-Ruiz.

## Materials Availability

This study did not generate new unique reagents.

## Data and Code Availability

- Raw electrophysiological and mice behavioral data have been deposited on servers at Cornell University, and are available upon request by contacting the lead contact.
- The original code that supports the findings of this study can be downloaded from Github (https://github.com/ayalab1/neurocode).
- Any additional information required to reanalyze the data reported in this paper is available from the lead contact upon request.

## Acknowledgments

The authors thank members of the Oliva and Fernandez-Ruiz labs for providing useful feedback on the manuscript. This work was supported by NIH grant R00MH122582 and R01MH130367 (AO); NIH grant R01MH136355, DP2MH136496, Sloan Fellowship, Whitehall Research Grant, Klingenstein-Simons Fellowship and Pershing Square Foundation’s MIND Prize (AFR); NIH grant F32MH134673 (H.L.R); Eric and Wendy and Schmidt AI in Science Postdoctoral Fellowship (R.T.).

## Author contributions

H.L.R., R.T., G.A.N., A.O. and A.F-R. conceived and designed experiments and analyses and wrote the manuscript. H.L.R., G.A.N., A.G., P.P., and R.T. performed experiments. R.T. and H.L.R. conducted the analysis.

## Declaration of interests

The authors declare no competing interests.

## STAR★METHODS

### Experimental model and study participant details

All experiments complied with guidelines established by the National Institutes of Health and approved by the Cornell University Institutional Animal Care and Use Committee (IACUC). All mice were housed in vivariums on a 12-hour light/dark cycle with a maximum of 5 animals per cage with *ad libitum* access to water and food. After surgery, mice were individually housed. Adult C57BL/6J (The Jackson Laboratory #000664, ∼25-35g, ∼3-6 months) or adult ‘hybrid’ offspring (∼35-45g, ∼3-6 months) were used for experiments not involving optogenetic manipulation. Hybrid mice were the first-generation offspring from a cross between C57BL/6J and FVB/NJ (The Jackson Laboratory, #001800)^67^. For optogenetic manipulation, transgenic animals or viral approaches (see Viral Injections) were used to express Channelrhodopsin (ChR2/H134R) in pyramidal neurons. For the transgenic approach, offspring of Ai32 (The Jackson Laboratory, #024109) and CamKIIɑ-Cre homozygous animals (The Jackson Laboratory, #005359) were used to express ChR2/H134R in pyramidal neurons.

## Method details

### Surgical Procedures

For silicon probes and viral injections, mice were anesthetized with isoflurane, and coordinates were taken following stereotaxic techniques following previously reported methods ^23,36,46^. Briefly, fur was removed using Nair and skin was washed three times with alcohol and iodine. A skin incision was then made and craniotomies were performed. For hippocampal virus injections, craniotomies were made at AP -2.3 mm, ML ±1.6 mm, DV -1.1 mm from Bregma, and AP -2.7 mm, ML ±2.5 mm, DV -1.25 mm. For CA1 silicon probe implantation, craniotomies were made at AP -2.5 mm, ML -2.0 mm. For the PFC, probes were inserted in a craniotomy made at AP +1.85 mm, ML ±0.35 mm, DV 1.5 mm. Silicon probes (128-channels probes from Diagnostic Biochips or 64-channel probes from Neuronexus) were attached to micro-drives (Cambridge Neurotech or custom-printed drives). For optogenetics, 200 𝜇m optical fibers (ThorLabs, part# FP200ERT) were glued to a probe. A stainless-steel screw (JI Morris, 000-120 x 1/16) attached to a thin wire (Phoenix Wire Inc., cat# 36744MHW) was used to connect the ground screw over the cerebellum. Next, custom 3D printed baseplates with copper mesh attached were glued to the skull using metabond (C&B Metabond Quick Adhesive Cement System). Probes were inserted above the target region, and the base of microdrives was secured to the skull using dental cement (UNIFAST Trad Acrylic Resin). Craniotomies were covered with Dura-gel (Cambridge NeuroTech Dura-Gel). Ground wires were connected to probes’ ground wires and finally to the shielding copper mesh. Animals were allowed to recover for a week before recording started, and they were given carprofen subcutaneously for the first 3 days. After recovery, probes were slowly moved between 50 𝜇m to 200 𝜇m steps per day until the desired location was reached, where LFP physiological landmarks and characteristic LFP patterns were used to identify hippocampal and cortical layers^46,68^.

### Viral Injections

Stock AAVs were acquired from Addgene. AAV-CamKIIɑ-ChR2/H13R-EYFP (3.4 x 10^13^ gc/ml, Addgene #26969-AAV5) was used. Specifically, 400nl of AAV-CamKIIɑ-ChR2/H13R-EYFP full virus titer was injected into the hippocampus bilaterally (see Surgical Procedures). Injections were delivered through a pulled pipette using Nanoject III (Drummond Scientific Company, 3-000-207). Animals were allowed two weeks for recovery and for viral expression to take effect before probe implantation (see Surgical Procedures). The pAAV-CaMKIIa-hChR2(H134R)-EYFP was a gift from Karl Deisseroth (Addgene plasmid # 26969; http://n2t.net/addgene:26969; RRID: Addgene_26969)^69^.

### Recording System and Data Preprocessing

Recordings were conducted with Intan RHD USB interface board or Intan Recording System. The sampling rate was 20,000 Hz, and amplification and digitization were done in the head stage by Intan RHD Electrophysiology Amplifier Chips. Acquisition was done with RHX Data Acquisition Software. LFPs were down-sampled to 1250 Hz before analysis. For video recording, an overhead mounted Basler Ace camera (Basler, acA3100-200uc) was used to record video simultaneously with neural signals. The camera’s frame rate (40 Hz) was recorded as digital inputs to Intan recording systems. Mice were tracked with custom DeepLabCut^70^ models trained on recorded videos and tracking was synchronized with individual frame timestamps.

### Behavioral Tasks

Before surgery, animals were exposed to an open field environment to decrease neophobia after implantation. After surgery and recovery, animals were handled daily and habituated to the experimenter, room, open field environment, and recording cables for at least a week before the start of behavioral experiments. Recordings of brain signals were done during the object displacement memory task, as previously described ^46^. Specifically, ∼2 hours of pre-sleep was recorded before a 5- minute open field exploration. After open-field habituation, mice were trained to explore two identical objects for 5-minute periods. Depending on the experiment, animals were allowed a single 5-minute exploration of objects or three 5-minute exposures to objects. After object training, animals were allowed to sleep either 1 or 4 hours post-sleep, depending on experimental conditions. After post-sleep, animals were tested in a 5-minute single trial where one of the identical objects was moved to a new location. A discrimination index was used to assess memory performance ^71^. Specifically, the discrimination index is calculated as:

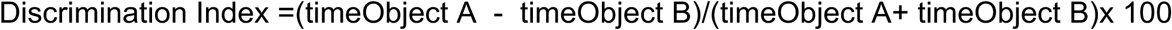

time Object A is the time during the test exploring the unmoved object, and time Object B is the time during the test exploring the moved object. The discrimination index results in a score between 1 and -1, where 1 corresponds to time spent studying only the moved object, -1 corresponds to exploring the unmoved object. Only sessions where animals explored for at least 5 seconds were used for behavioral experiments ^46^.

### Optogenetic manipulations and real-time SWR Detection

During the first 2-hours of post-task sleep, real-time SWR detection was conducted as done previously ^21,23^. Three LFP channels were used for online SWR detection. An LFP channel with the strongest ripple amplitude was selected for the ripple channel, filtering between 100 and 300 Hz. Another channel with the largest sharp-wave amplitude was selected to detect sharp-wave, filtering between 10 and 40 Hz. A third noise channel was selected by choosing a channel in the neocortex where no SWR activity was present. The noise channel was filtered between 100 and 300 Hz. The root-mean-square (RMS) of the three signals was then computed by a custom-made diode bridge rectifier (1N4001 diodes, 560 kOhm resistor, 68 nF capacitor). The filtered and rectified LFP were sent to a programmable real-time processor (Cambridge Electronic Design, CED) at 20,000 Hz sampling rate. Custom scripts in SpikeDetect2 (CED) were used to manually set the detection thresholds for ripple, sharp-wave, and noise for each animal individually. SWRs were defined as events crossing the ripple and sharp-wave thresholds but not crossing the noise channel threshold.

Once a SWR was detected, a small amplitude, 100 ms, light-pulse was delivered through bilateral optic fibers. On average, the delay between SWR detection and optogenetic pulse delivery was 22.9 ± 0.2 ms (n = 10,727 events). The light intensity was manually adjusted for each animal on the day of the experiment while the animal was in its homecage, where light power was increased until a ripple was produced. A trapezoid pulse with a 20 ms initial ramp and 20 ms ramp after the pulse was used ^21,72^. As a control, in different sessions, SWR were detected during post-task sleep and the same light-pulse was delivered after a random delay between 500 and 1,000ms (i.e. “Delayed stimulation” see Figure S8D-E^21,23^).

### Tissue Processing and Immunohistochemistry

After experiments, mice were deeply anesthetized under high-concentration (4-5%) isoflurane anesthesia. Mice were then perfused transcardially with 0.9% phosphate buffer saline (PBS) solution, after which 4% paraformaldehyde (PFA) solution was perfused. Brains were kept for 24 hours in PFA before sectioning them into 70 𝜇m thick slices^73,74^. Slices were then stained with DAPI (Thermo Fisher, #D21490) and incubated for 30 minutes before washing. After washing, slices were mounted on glass slides with a Polyvinyl alcohol mounting medium with DABCO antifading (Millipore Sigma, #10981). Slides were imaged with a confocal microscope (Zeiss LSM 800 - acquired through Cornell University Biotechnology Resource Center, with NYSTEM (CO29155) and NIH (S10OD018516)) to image DAPI and virally expressed EYFP (see Virus Injections).

## Quantification and statistical analyses

### Spike Sorting and Single Unit Classification

Spike sorting was performed semi-automatically using Kilosort^75^ (https://github.com/cortex-lab/KiloSort). After sorting, manual curation was done in Phy (https://github.com/kwikteam/phy-data) using custom-designed plugins to acquire single well-isolated units (https://github.com/petersenpeter/phy1-plugins). Clusters were manually inspected for waveforms, auto-correlograms, and isolation distance metrics. Multi-units, noise clusters, or poorly clustered units were discarded, whereas well-isolated units were classified into putative cell types using the CellExplorer Matlab package (https://cellexplorer.org/)^76^. Three cell types were assigned: pyramidal, narrow-waveform interneurons, and wide-waveform interneurons. Two key metrics separated different putative cell types: the burst index and trough-to-peak latency ^77^. The burst index was calculated as the average number of spikes in the 3-5 ms bins of a cell’s autocorrelogram divided by the average number of spikes in the 200 to 300 ms bins. The average waveform from the recording site with maximum amplitude was taken to determine the trough-to-peak latency. Only well-isolated neurons with at least 100 spikes were kept for further analysis.

### Sleep and Brain States Classification

Sleep and brain state scoring was done as previously described^29,46,78,79^. State scoring used broadband LFP, narrow band theta frequency, and estimated electromyogram (EMG) to differentiate brain states. From broadband LFP, spectrograms were calculated using a fast Fourier transform with 10 s epochs and a sliding window of 1 s. Principal Component Analysis was then computed from the Z-transform of the spectrogram from 1 to 100 Hz. The first component reflected the power in the low-frequency range (<20 Hz) with opposite weighted power at higher frequencies (>32 Hz) and distinguished non-rapid eye movement (NREM) from non-NREM periods ^78^. For theta epochs, a channel with high theta power was selected, and theta dominance was quantified as the ratio of power in the 5-10 Hz and 2-16 Hz frequency bands ^79^. For EMG, the zero-lag correlation of filtered signal between 300-600 Hz (Butterworth filter with filter padding between 275-625 Hz) between recording sites was used to estimate EMG activity. Pairwise correlations between channels of non-adjacent probe shank were calculated, and the mean of all correlations in time bins of 0.5 s was used as an EMG score ^80^. Soft sticky thresholds were used on broadband LFP, narrow band theta, and EMG were used to determine states. High LFP in the first principal component of broadband spectrogram and high EMG were considered NREM, high narrow band theta activity and low EMG were considered rapid eye movement (REM), and remaining data (i.e., high EMG) were taken as waking state.

To prevent variations in the total amount of recorded NREM sleep from affecting our results given that SWR rate and reactivation are known to change throughout NREM sleep, we defined “baseline sleep” as the first cumulative hour of NREM sleep in the rest session recorded before training, and “post-task sleep” as the first cumulative hour of NREM sleep in the rest session recorded after training.

### SWR analyses

For offline SWR detection, the wide-band signal was band-pass filtered (difference-of-Gaussians; zero-lag, linear phase FIR), and instantaneous power was calculated by clipping at 4 SD, rectified, and low-pass filtered. The low-pass filter cut-off was at a frequency corresponding to p cycles of the mean bandpass (for 80-250 Hz band-pass, the low-pass was 55 Hz). One channel around the CA1 pyramidal layer was chosen for the ripple detection, and one channel from CA1 str. radiatum was chosen for sharp wave detection. Candidate events were identified as local minima in the sharp-wave band-filtered signal, and the corresponding ripple power was recorded. K-means clustering was used to define clustering of SWR from non-SWR events, and manual curation was used to better define the boundary and remove outliers. The events were then expanded until the (non-clipped) ripple power fell below 1 SD and the sharp-wave band-filtered signal fell below 0.5 SD above the local (±5 s) median; short events (<15 ms) and longer events (>400 ms) were discarded.

LFP signals were extracted around detected ripple events. For each event, a wavelet spectrogram was computed on the detrended LFP segment within ±500 ms of the ripple peak. The mean wavelet spectrogram for SWRs in baseline sleep and SWRs in post-task sleep, and each was corrected by subtracting power at -200 ms. Subsequently, the minimum and maximum power within the 150 Hz frequency band during the ±100 ms window around the event were used to normalize the spectrogram as w = (w-min)/(max-min).

To compare SWR spectral power in the ripple band (e.g. Figure 1E), the power values in the ripple band (150−250 Hz) within ±50 ms around the ripple peak from the baseline sleep spectrogram and the post-task sleep spectrogram for each session were normalized together by subtracting the combined mean of both and dividing by their combined standard deviation. A Monte-Carlo test was then used to compare the normalized baseline sleep power to the normalized post-task sleep power for each task condition.

To compare SWR spectral profiles (e.g. Figure 1F), the mean spectra within ±100 ms around the ripple peak from baseline sleep and post-task sleep for each session were normalized together by subtracting the combined mean of both conditions and dividing by their combined standard deviation. A Monte-Carlo test was then used to compare the normalized baseline sleep spectra to the normalized post-task sleep spectra for each task condition.

SWR start and end are defined as part of the offline SWR detection described above. SWR peak was defined as the moment of maximal ripple amplitude power within the event. SWR asymmetry was defined as how far the ripple peak deviated from the center of the event, computed as the absolute deviation of the peak from the midpoint between ripple start and end: |(peak - start) / (end - start) - 0.5|, with values near 0 indicating a symmetrical event. SWR frequency was defined as the frequency at which the wavelet spectrogram reached its maximum within ±100 ms of the ripple peak, and SWR amplitude was the corresponding power at that frequency. To visualize the SWR features, we included SWR duration, SWR amplitude, SWR frequency, SWR asymmetry, and reactivation strength of PFC cell assemblies (see below), and reduced the dimensions using an unsupervised Uniform Manifold Approximation and Projection (UMAP)^36,81^ and plotted the first two dimensions, color-coding by feature.

To select SWRs with respect to their size in terms of both duration and amplitude, we designed a compound measure combining these features. For each session, SWR amplitude and SWR duration were z-scored using the mean and standard deviation of the SWR amplitude and SWR duration values, respectively, measured from baseline sleep SWRs. SWR size was defined as the minimum of these two values, and events with an SWR size exceeding the mean size of baseline sleep SWRs were classified as “large SWRs,” and other SWRs were classified as “small SWRs”.

### LFP coherence analyses

Coherence was measured between LFP signals in the PFC (pre-limbic cortex) and CA1 pyramidal layer. LFP epochs recorded during movement periods in the task were concatenated. Epochs that contained artifacts were rejected. Spectra were estimated by Fourier transform of the LFP. Coherence was calculated as the magnitude of the summed cross-spectral density between two LFP time series, normalized by respective power spectra.

### Current Source Density analysis

To address the contribution of different synaptic inputs to the LFPs recorded in the CA1 subregion, we employed current source density (CSD) analysis^36,42,82^(Fig. 1L). First, event-triggered LFP signals were isolated for small and large SWR separately. For each of them, CSD analyses were applied to the profile of the CA1 LFPs. The different layers (i.e., stratum oriens – ori.-, radiatum – rad.- and lacunosum-moleculare -l-m.-) associated with the main inputs into CA1 (i.e., CA2, CA3 and entorhinal inputs respectively) were identified by the voltage reversal of LFP signals and depth profile of currents sources and sinks (from CSD analysis), assuming current sources as positive CSD values and current sinks as negative CSD values as described previously^36,42,82^ ^rf^. The strength of each input was estimated as the power of the CSD in their corresponding layers. Specifically, the CSD value for each input (ori., rad. or l-m.) was calculated as the absolute magnitude of the corresponding sinks in each layer. Then, a *ratio* was calculated as the CSD magnitude during SWRs as:

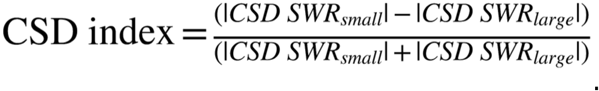

### Cell assembly analyses

Cell assemblies were detected using an unsupervised statistical framework based on a hybrid principal component analysis (PCA) followed by independent component analysis (ICA) as reported previously^24,41,64^. In brief, for each region (the hippocampus and the prefrontal cortex), spike trains of each neuron were binned in 50-ms intervals for the whole session, the pairwise firing correlation coefficients were computed for all pairs. Next, we calculated the number of assemblies based on those principal components whose eigenvalues exceeded the threshold for random firing correlations (using the Marčenko-Pastur law), ensuring that each of the detected patterns explained more variance of the spike train correlation matrix than other patterns that would result from independently firing neurons. Independent component analysis (fast-ICA algorithm) was then used to determine for each assembly (component) the vector of weights with which each neuron’s firing contributes to that assembly. The strength of each assembly *i*’s activation for a given time bin *k* was computed as follows:

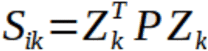

where Z is the population activity matrix of the z-scored firing rate of each unit, and *P* is the outer product of the component *i*’s weights, in which the diagonal has been set to zero so that isolated spikes from individual units do not contribute to *S*.

The goal of the cell assembly analysis was to study the patterns associated with the learning experience (i.e. object exploration). However, because the task design required the animals to be habituated to the environment in advance, we expected a number of cell assemblies to have formed during this habituation period, reflecting task-related variables (such as the environment layout, distal cues, etc.) that are orthogonal to the learning experience. We therefore discarded cell assemblies with significant reactivation in baseline sleep before object exploration from subsequent analyses. Unless otherwise specified, throughout the text, the term ‘assembly reactivation’ refers to the reactivation in SWRs of cell assemblies identified using this PCA-ICA based approach.

To detect cross-structural cell assemblies, we followed a procedure described previously ^29,47^. Briefly, we combined the recordings in the hippocampus and the prefrontal cortex and performed assembly detection as described above. For a given assembly, the weight vector was normalized to have unit Euclidean norm, and units whose weights exceeded 1/√N, where N is the number of recorded units, were considered assembly members. Cell assemblies with members in both the hippocampus and the prefrontal cortex were retained as cross-structural cell assemblies.

Assembly responses to SWRs were quantified as the mean reactivation strength within 0–100 ms of SWR onset. To allow pooling across assemblies, responses were z-scored within each assembly. For analyses requiring classification of individual SWR responses (Figure 1; Figure S2; Figure S7), responses exceeding z = 1.96 (P < 0.05) were considered significant; SWRs without PFC significant reactivation were considered as SWRs in which none of the detected PFC cell assemblies reached a significant response.

### Pairwise reactivation

Pairwise reactivation analyses were performed as described previously ^24,83^. Briefly, we binned the activity of all units during SWRs in baseline sleep, and used this spike count matrix to compute a pairwise cross-correlation matrix C_baseline_. Similarly, we binned the activity of all units in 100 ms during movement intervals of training of the object task, and used this spike count matrix to compute a pairwise cross-correlation matrix C_behavior_. Finally, we binned the activity of all units during SWRs in post-task sleep, and used this spike count matrix to compute a pairwise cross-correlation matrix C_post_. We used linear regression between C_baseline_ and C_behavior_ and between C_baseline_ and C_post_ to remove the portion of the variance that can be explained by preexisting patterns. The reactivation of pairwise behavioral patterns in post-task sleep was quantified as the correlation between the resultant residuals R_behavior_ and R_post_. This method was used in Figure 3G,H, and Figure 4A.

### Hierarchical bootstrap

Hierarchical bootstrap ^84^ was performed to analyze data with hierarchical structure. Briefly, bootstrap datasets were created by resampling with replacement following levels of hierarchical order in the order of animals followed by sessions followed by the within-session observations. Each resampled bootstrap dataset was used to calculate an estimate of the mean, in a total of 5000 resampling times.

For the pairwise reactivation analysis, hierarchical bootstrap was used to estimate a distribution of correlation values between R_behavior_ and R_post_ . To this end, R_behavior_ and R_post_ were resampled together according to the hierarchical structure, and the correlation between them was computed for each bootstrap sample.

In all cases (whether estimating a mean or a correlation), the P-value reflects the direct comparison of the distribution of estimates to zero, or comparisons of different distributions to each other, after controlling for the other nesting variables.

### Statistical Analysis

Data analysis was performed using custom routines in Python and MATLAB (MathWorks). No specific analysis was used to estimate minimal population sample or group size, but the number of animals, sessions and recorded cells were larger or similar to those employed in previous related work ^20,25,36,46^. Unless otherwise noted, non-parametric Wilcoxon rank-sum or Wilcoxon signed-rank test was used for unpaired and paired data comparisons respectively, and ANOVA and Kruskal-Wallis was used for multiple comparisons. All statistical tests were two-tailed with *p* < 0.05 as the cutoff for statistical significance, which is indicated by asterisks (\**p*<0.05, \*\**p*<0.01, \*\*\**p*<0.001, and \*\*\*\**p*<0.0001). Error bars show SEMs, and boxplots show median (central mark), 75th (box), and 90th (whiskers) percentile, unless indicated otherwise. Due to experimental constraints of optogenetic experiments, experimenters were not blind to these manipulations.

---

**Figure S1:**
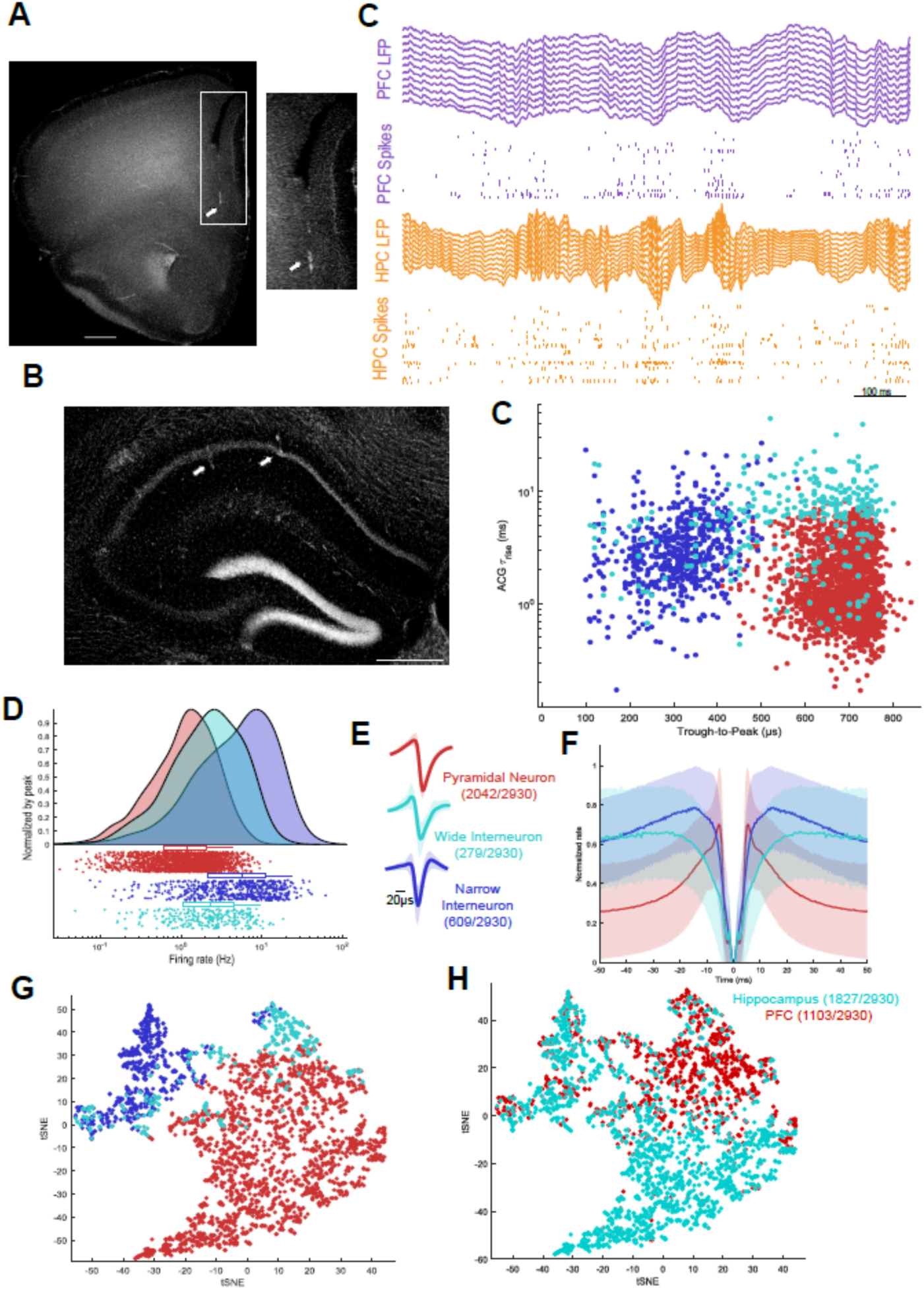
Hippocampus and PFC histology and dual recordings. Related to. Figure 1. A) Histology of Prefrontal Cortex (PFC) with DAPI staining with (left) and inset of white box (right). Arrows indicate fiber tracks (scale bar 500um). **B**) Histology of hippocampus with DAPI staining. Arrows indicate fiber tracks (scale bar 500um). **C**) Example PFC-CA1 recordings (only one shank from each probe is shown): PFC LFP (top, purple), PFC spikes (middle top, purple), CA1 LFP (middle bottom, gold), and CA1 spikes (bottom, gold). **D**) Scatter plot for all recorded units using autocorrelogram (ACG) tau-rise (ms) versus Trough-to-Peak (us) of Pyramidal Neurons (red, n = 2042), Wide Interneurons (cyan, n = 279) and Narrow Interneurons (blue, n = 609). **E**) Top, average waveforms of Pyramidal Neurons, Wide Interneurons and Narrow Interneurons . Bottom, normalized ACG of Pyramidal Neurons (red), Wide Interneurons (cyan), and Narrow Interneurons (blue). **F**) Average Firing rate (Hz) of Pyramidal Neurons (red), Wide Interneurons (cyan) and Narrow Interneurons (blue). **G**) t-distributed stochastic neighbor embedding (tSNE) of waveform and firing rate characteristics of Pyramidal Neurons (red), Wide Interneurons (cyan), and Narrow Interneurons (blue). **H**) tSNE using the same waveform and firing rate characteristics for PFC (red, n = 1103) and Hippocampus (cyan, n = 1827) units from 21 animals.

**Figure S2:**
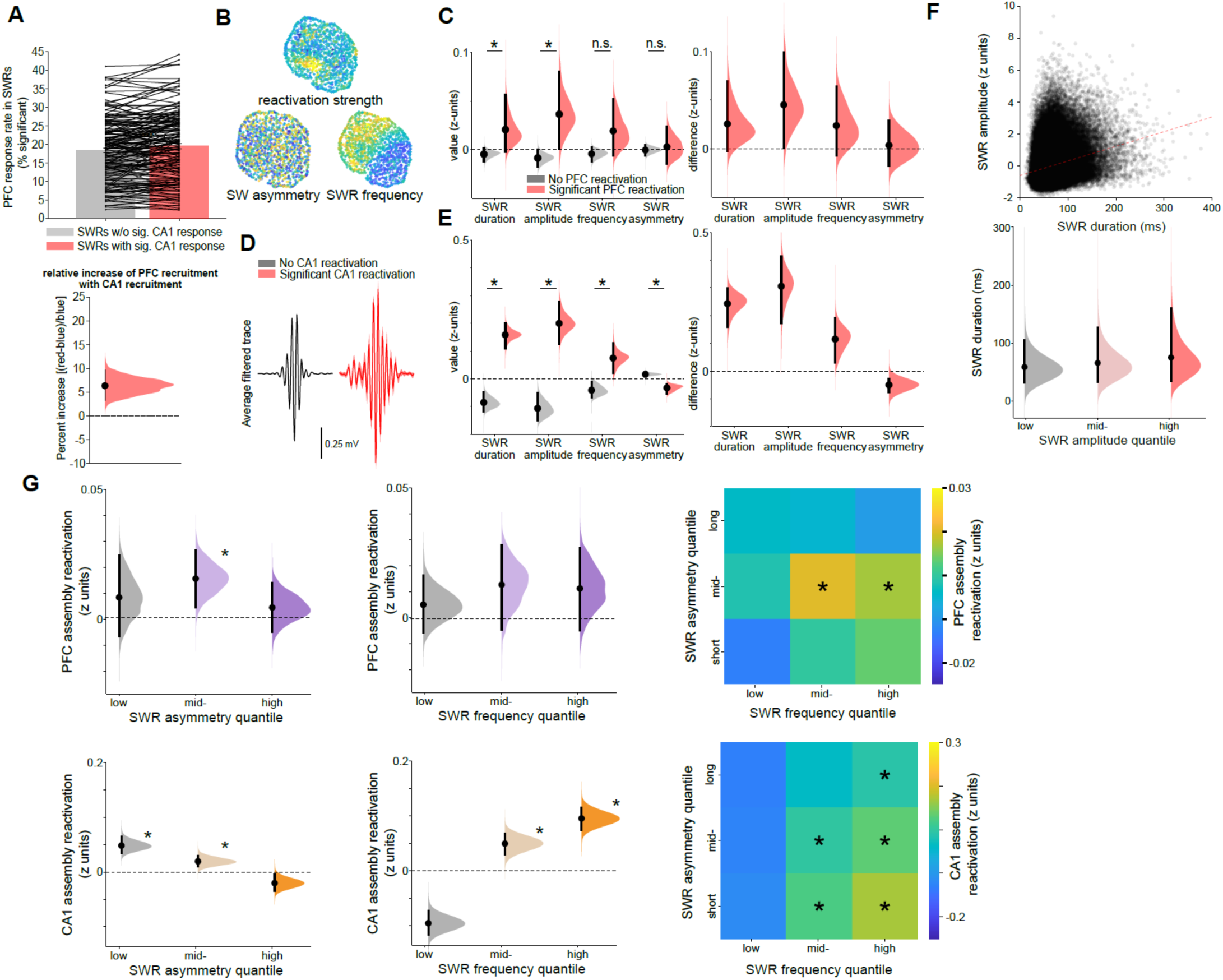
PFC and CA1 reactivation and other SWR properties. Related to Figure 1. **A**) PFC reactivation increased in SWRs with CA1 reactivation. Top: probability of PFC reactivation following SWRs with (red) and without (grey) significant CA1 reactivation (P = 0.00067, N = 175 sessions; Wilcoxon signed rank test). Bottom: hierarchically bootstrapped estimate of the percent increase in PFC reactivation probability that is explained by CA1 reactivation (P = 0). **B**) Additional SWR properties represented in the same UMAP space as in Figure 1. **C**) Left: same as Figure 1E for additional SWR properties. Right: Hierarchically bootstrapped difference in normalized SWR properties between SWRs associated with and without significant PFC assembly reactivation. Note that duration and amplitude, but not frequency or asymmetry were associated with PFC reactivation (P = 0.048 /0.025 /0.092 /0.43 for SWR duration, amplitude, frequency, and asymmetry, respectively; hierarchical bootstrap test). **D**) Same as Figure 1D but for CA1 reactivation. **E**) Same as B but dividing SWRs depending on the reactivation of CA1 cell assemblies. Unlike PFC reactivation, CA1 reactivation correlated with all SWR features (P = 0 / 0 / 0.0008 / 0.0012 for SWR duration, amplitude, frequency, and asymmetry, respectively). **F**) Top: scatter plot between SWR duration and normalized amplitude. There was a moderate correlation between duration and amplitude (red line, Pearson’s r = 0.23, P < 0.001) but many events had high amplitude and short duration and vice versa. Bottom: hierarchically bootstrapped estimates of the mean SWR duration classified by SWR amplitude were similar. **G**) Upper panels are the same as Figure 1J but for SWR frequency and asymmetry. Unlike duration and amplitude, no clear relationship could be found between PFC reactivation and SWR asymmetry (left: P = 0.16 / 0.006 / 0.22 for low-, mid- and high-asymmetry SWRs, respectively; hierarchical bootstrap test) or frequency in upper middle panel (middle: P = 0.22 / 0.074 / 0.102 for low-, mid- and high-frequency SWRs, respectively). Upper right: PFC assembly responses to SWR ripple quantile grid split according to both asymmetry (y-axis) and frequency (x-axis) (*P < 0.05, Wilcoxon signed-rank tests). Same as above but for CA1 reactivation. Bottom left is CA1 reactivation correlated with SWR asymmetry (left: P = 0 / 0 / 0.99 for low-, mid- and high-asymmetry SWRs, respectively; hierarchical bootstrap test) and bottom middle is frequency (middle: P = 1 / 0 / 0 for low-, mid- and high-frequency SWRs).Bottom right: HPC assembly responses to SWR ripple quantile grid split according to both asymmetry (y-axis) and frequency (x-axis) (*P < 0.05, Wilcoxon signed-rank tests).

**Figure S3.**
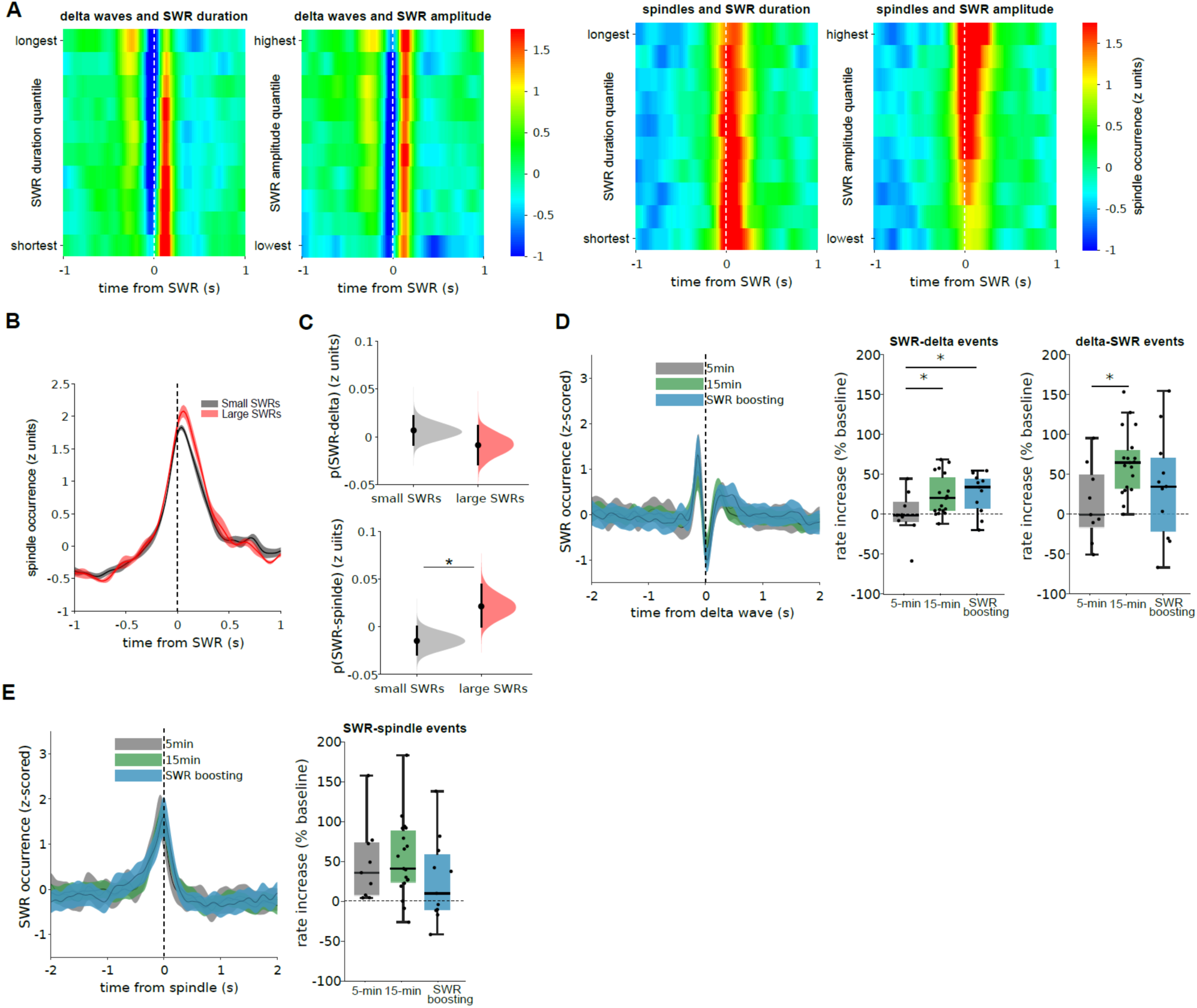
Coupling of SWRs with cortical delta waves and spindles.Related to Figure 1 and 3 **A)** Peri-event time histograms showing the occurrence (color) of cortical delta waves (left) and sleep spindles (right) centered on SWRs. SWRs are grouped in quantiles as a function of duration (left) and ripple amplitude (right). **B)** Mean peri-event time histogram ofspindle occurrence around small (black) and large SWRs (red). Mean ± SEM. **C)** Top: probability of a delta wave occurring after SWRs (50 to 200 ms) was similar for small and large SWRs (P = 0.78, N=18 sessions in 10 animals); bottom: probability of spindle occurrence (-400 to -100 ms) was higher for large SWRs than small SWRs (P = 0.024, hierarchical bootstrap test; N=18 sessions). **D)** Left: Averaged peri-delta wave histograms showing the occurrence of SWRs displayed a similar profile in post-task sleep for all conditions. Middle: increase of SWR-delta events (with the delta wave following the SWR from 50 to 200 ms) in post-task sleep relative to baseline sleep was higher following 15 minute training than 5 minute training (P = 0.012, Wilcoxon ranksum test; N=19 /9 sessions, respectively), but it was similar between SWR boosting and 5 minute training (P=0.028, N=11 /9). Right: increase of delta-SWR events (with the delta wave preceding the SWR from -400 to -100 ms) in post-task sleep relative to baseline sleep was higher following 15 minute training than 5 minute training (P = 0.006, N=19/ 9), but it was similar between SWR boosting and 5 minute training (P=0.22, N=11/ 9 sessions, respectively). **E)** Left: Averaged peri-spindle histograms showing the occurrence of SWRs displayed a similar profile in post-task sleep for all conditions. Right: increase of SWR-spindle events in post-task sleep relative to baseline sleep was similar following 15 minute training as compared 5 minute training (P = 0.31, Wilcoxon ranksum test; N=19 /9), and during SWR boosting as compared to 5 minute training (P=0.94, N=11 /9).

**Figure S4.**
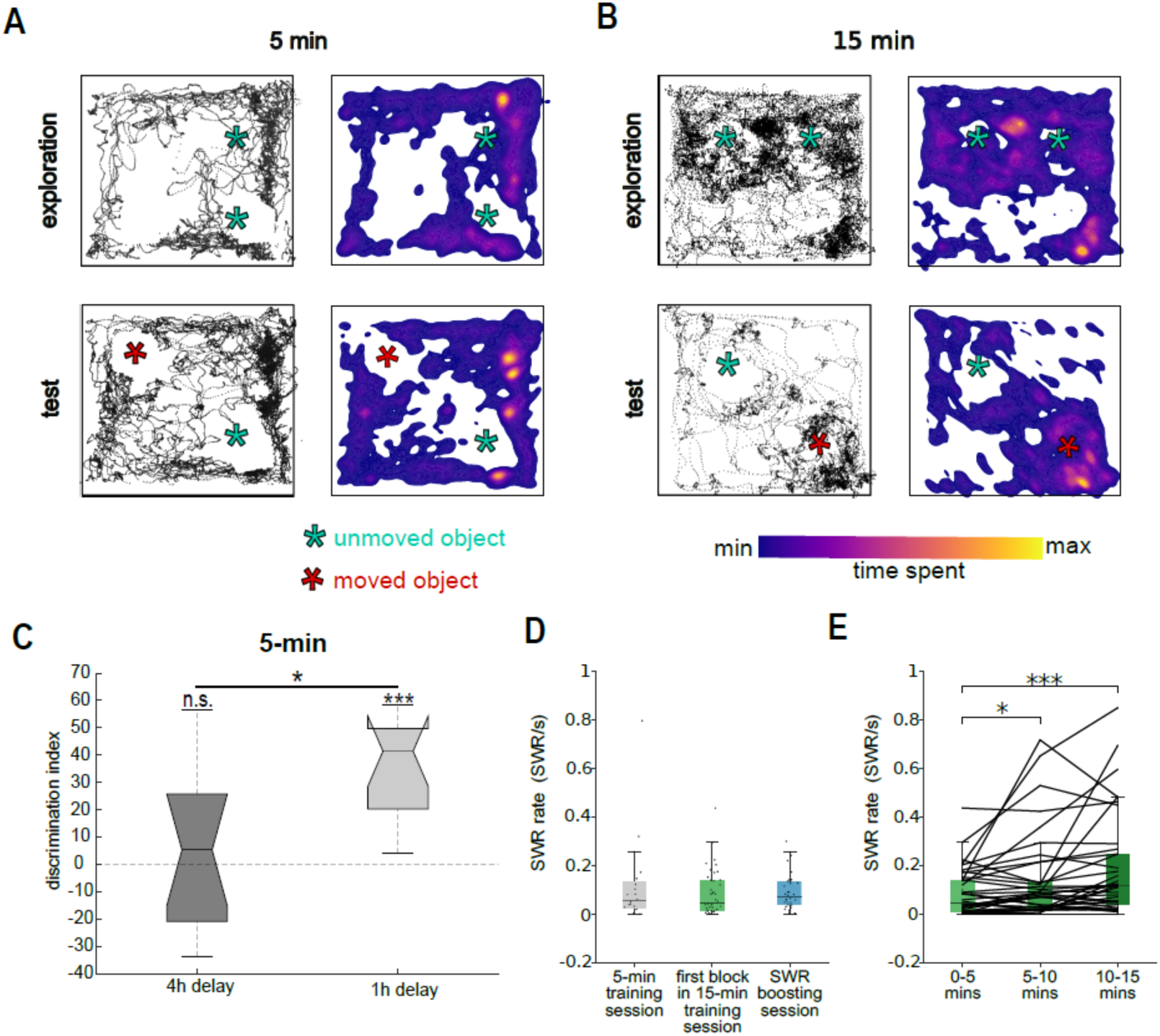
Examples of behavioral trajectories during the task. Related to Figure 2. **A**) Top: Animal movement (left) and occupancy (right, heatmap) for a representative 5-minute training session (top) and subsequent test (bottom). Cyan asterisks show the center of objects in the maze. Red asterisk represents moved object during test session. **B**) Same as A) but for a representative 15- minute training session and its test session. Note increased occupancy around displaced object during the test. **C**) Memory recall performance (object discrimination index) for 4 hours post-training (5- minute) sessions (same data as Figure 2B) and 1 hour post-training (5-minute) sessions (n = 13 mice, P = 2.4e-4, Wilcoxon signed rank test. 1 h vs 4 h test P = 0.01, Wilcoxon rank sum test). **D**) SWR rates during immobility periods in training sessions. SWR rates were not different between non SWR boosted sessions (grey and green) and sessions followed by SWR boosting (left: N = 23/34 sessions, P = 0.3123, Wilcoxon rank-sum test). **E**) In 15-minute training session, the awake ripple rate within the first 5-minute block was similar to 5-minute training sessions (left: N = 40/23 sessions, Wilcoxon rank-sum test), but it was increased in subsequent blocks (block 2 vs block 1, N = 40 sessions, P = 0.0109, Wilcoxon signed rank test; block 3 vs block 1: N = 40 sessions, P = 8.2e-6, Wilcoxon signed rank test).

**Figure S5.**
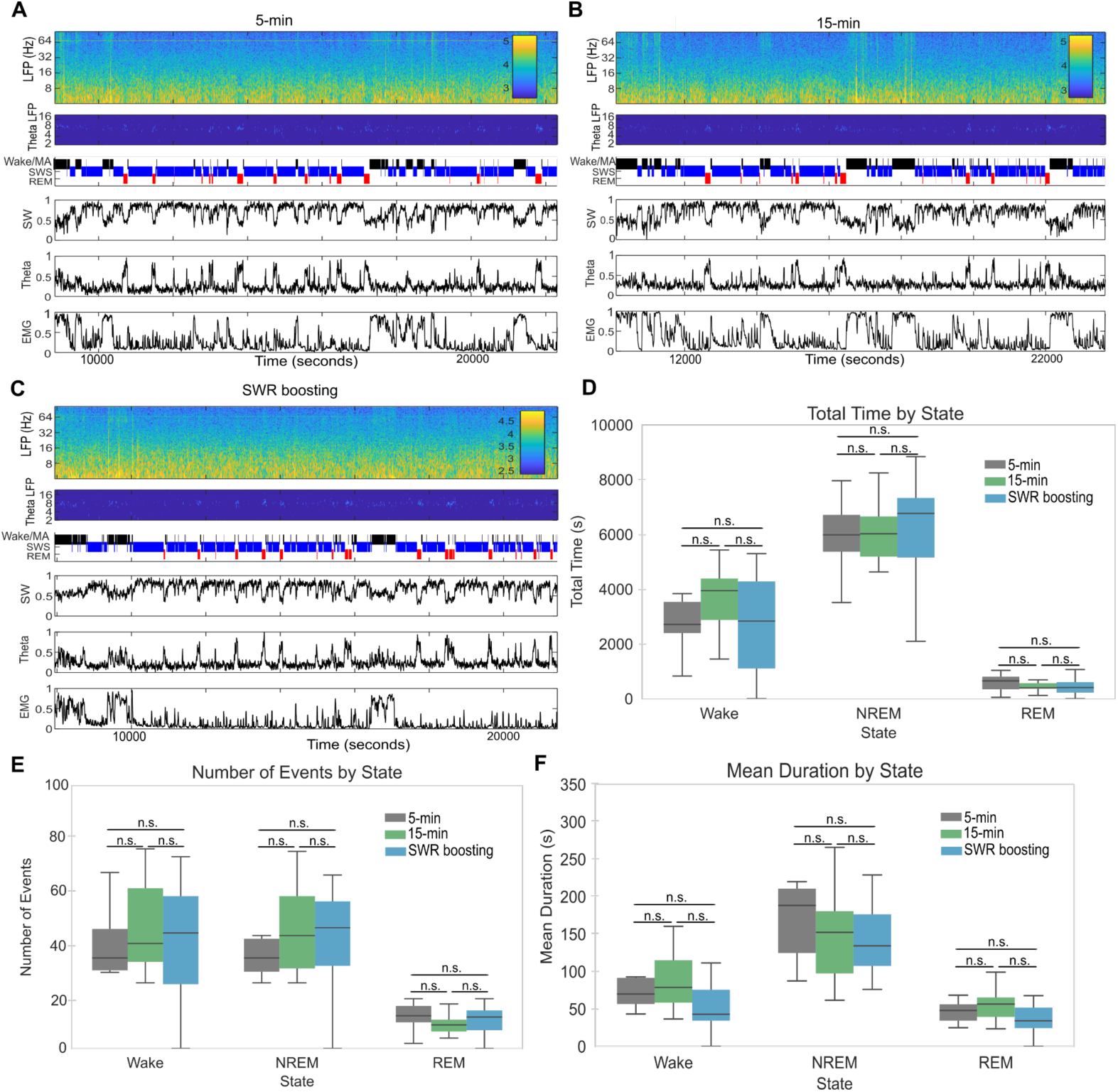
Sleep properties for 5-minute,15-minute conditions, and SWR Boosting. Related to Figure 2 and Figure 3. **A**) Representative example of sleep state scoring after a 5-minute training session. From top to bottom: LFP spectrogram, theta frequency power, brain state labels (Wake black, Slow Wave Sleep blue and REM sleep red), measure of Slow Wave sleep (SW, projected first PCA component on z-scored 1 to 100 Hz spectrogram), measure of Theta (ratio of powers in 5-10 Hz and 2-16 Hz), and measure of EMG (zero lag correlation in 300-600 Hz band between recording sites). **B**) same as A) but for 15-minute training session. **C**) Same as **B**) and **C**) but for SWR Boosting. **D**) Total time spent in each state during post-task from 21 animals for 5 minutes (grey), 15 minutes (blue), and SWR Boosting (orange): Wake (left, t-test, 5 minutes vs 15 minutes p = 0.12, 5 minutes vs SWR Boosting p= 0.08, 5 minutes vs SWR Boosting p = 0.84), NREM (middle, t-test, 5 minutes vs 15 minutes p = 0.49, 5 minutes vs SWR Boosting p = 0.64, 5 minutes vs SWR Boosting p = 0.96), and REM (right, t-test, 5 minutes vs 15 minutes p = 0.29, 5 minutes vs SWR Boosting p = 0.67, 5 minutes vs SWR Boosting p = 0.85). **E**) Number of bouts of each state for 5 minutes (grey), 15 minutes (blue), and SWR Boosting (orange) in Wake (left, t-test, 5 minutes vs 15 minutes p = 0.36, 5 minutes vs SWR Boosting p = 0.67, 5 minutes vs SWR Boosting p = 0.58), NREM (middle, t-test, 5 minutes vs 15 minutes p = 0.24, 5 minutes vs SWR Boosting p = 0.48, 5 minutes vs SWR Boosting p = 0.57), and REM (right, t-test, 5 minutes vs 15 minutes p = 0.143, 5 minutes vs SWR Boosting p = 0.82, 5 minutes vs SWR Boosting p = 0.38). **F**) Duration of bouts of each state for 5 minutes (grey), 15 minutes (blue), and SWR Boosting (orange) in Wake (left, t-test, 5 minutes vs 15 minutes p = 0.92, 5 minutes vs SWR Boosting p = 0.40, 5 minutes vs SWR Boosting p = 0.26), NREM (middle, t-test, 5 minutes vs 15 minutes p = 0.18, 5 minutes vs SWR Boosting p = 0.21, 5 minutes vs SWR Boosting p = 0.87), and REM (right, t-test, 5 minutes vs 15 minutes p = 0.32, 5 minutes vs SWR Boosting p = 0.42, 5 minutes vs SWR Boosting p = 0.07).

**Figure S6.**
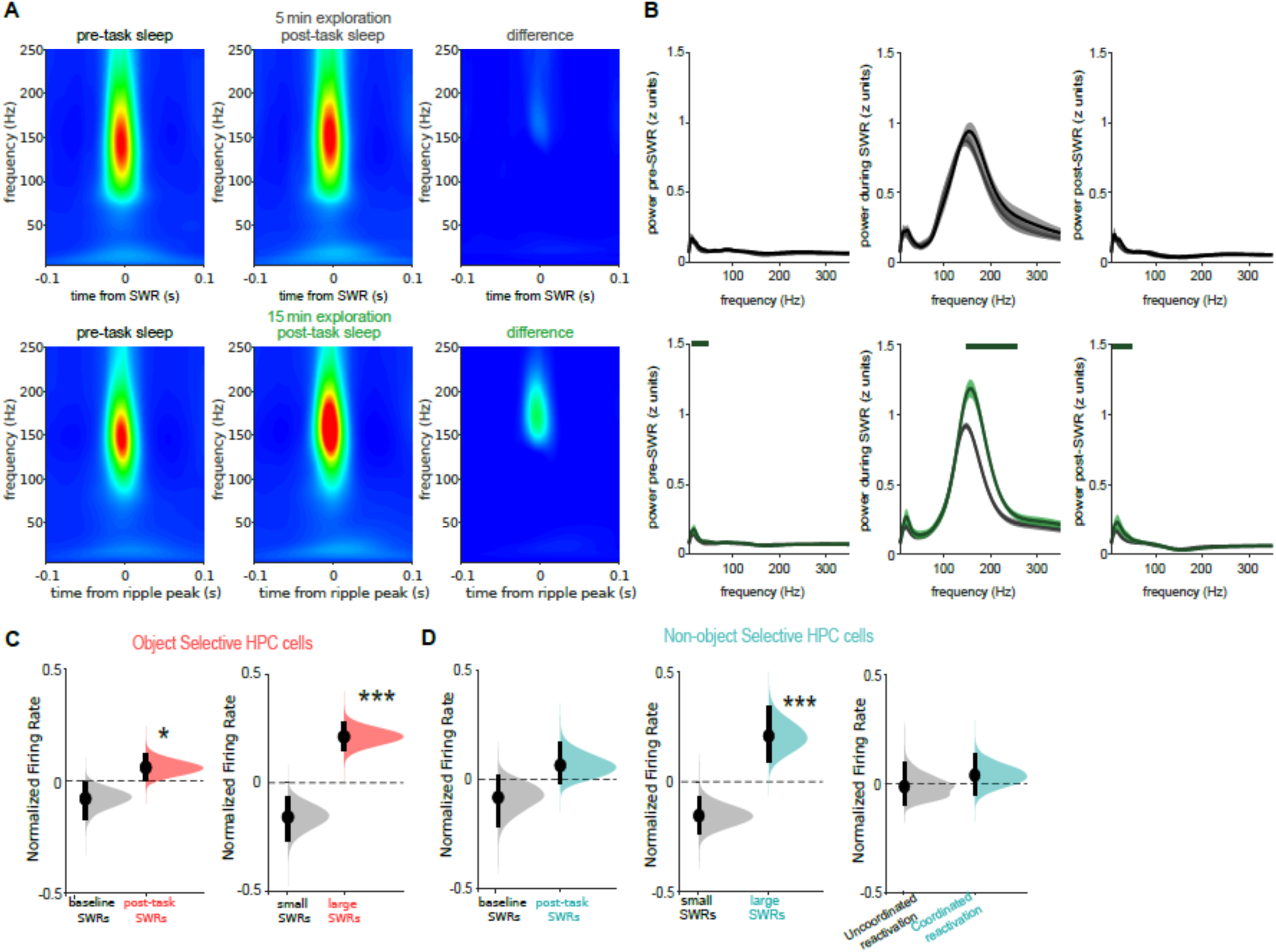
Additional SWR properties for different task conditions. Related to Figure 2 and 4 **A)** Average sleep SWR wavelet spectrograms extended to include lower frequencies for pre (left), post-task sleep (middle) and their difference (right), following either 5-minute (top, n = 20) or 15-minute (bottom, n = 38) exploration sessions. **B)** Spectral profile of the peri-SWR intervals: left: pre-SWR interval (75ms to 25ms before the SWR peak); middle: SWR interval (within 50 ms of the SWR peak); right: post-SWR interval (25 to 75ms after the ripple peak). Top: there were no significant changes in peri-SWR spectral properties in sleep after 5-minute exploration relative to baseline sleep (top; P>0.05, Monte Carlo test, n = 20 sessions), while the spectral profile was enhanced in the lower frequencies in the pre- and post-SWR intervals and in the ripple band in the SWR interval following 15-minute exploration relative to baseline sleep (bottom; P<0.05, n = 38 sessions; lines indicate the significant intervals). **C)** Normalized firing rate of spatially selective hippocampal cells firing preferentially around the objects in different SWRs. Left: firing rates in post-task SWRs were increased relative to firing rates in baseline SWRs (P = 0.019, hierarchical bootstrap test); Right: within post-task sleep SWRs, firing rates were higher in large SWRs relative to small SWRs (P = 0); **D)** Same as C but for spatially selective cells away from the objects. Left: spatially selective cells away from the object did not significantly increase their firing rates in post-task sleep relative to baseline sleep (P = 0.078). Middle: within post-task sleep SWRs, their firing rates were higher in large SWRs than small SWRs (P = 0). Right: within post-task sleep SWRs, their firing rates were similar in coordinated and uncoordinated CA1-PFC reactivation events (P = 0.21, hierarchical bootstrap test).

**Figure S7:**
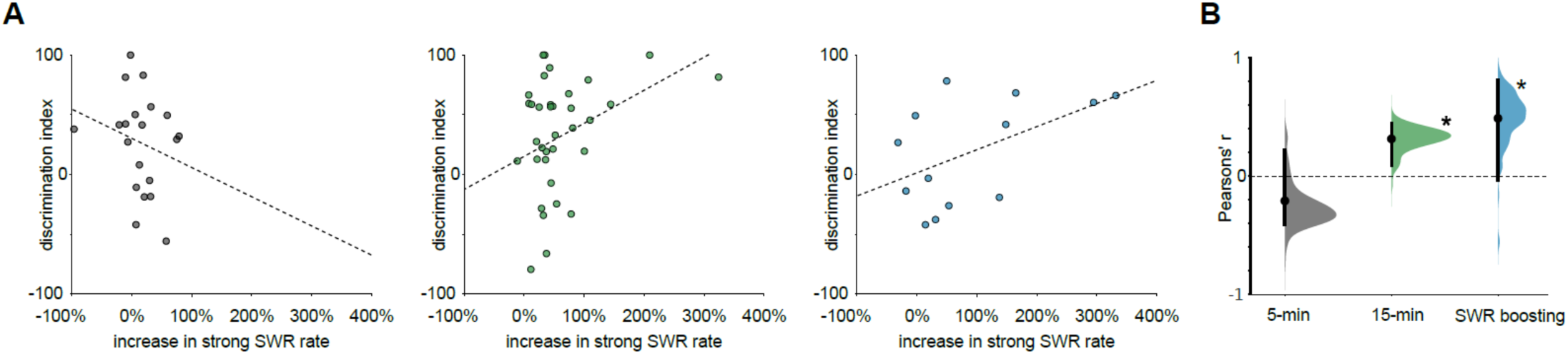
Increase in large SWR rate is associated with subsequent behavioral performance. Related to Figure 2 and 3. **A**) Correlation between large SWR rate increases relative to baseline sleep and memory recall performance (discrimination index) during test sessions. Left: sessions in which animals were trained for 5 min; middle: sessions in which animals were trained for 15 min; right: sessions in which animals were trained for 5 min and SWRs were subsequently optogenetically boosted. **B)** Hierarchical bootstrap estimate of the correlation (Pearson’s r) between large SWR rate increase and discrimination index for each session. Correlation was not significant following 5 min training (P = 0.915, hierarchical bootstrap test), but it was significant following for 15 training (P = 0.010, hierarchical bootstrap test), and following 5 training when SWRs were optogenetically boosted (P = 0.025, hierarchical bootstrap test).

**Figure S8:**
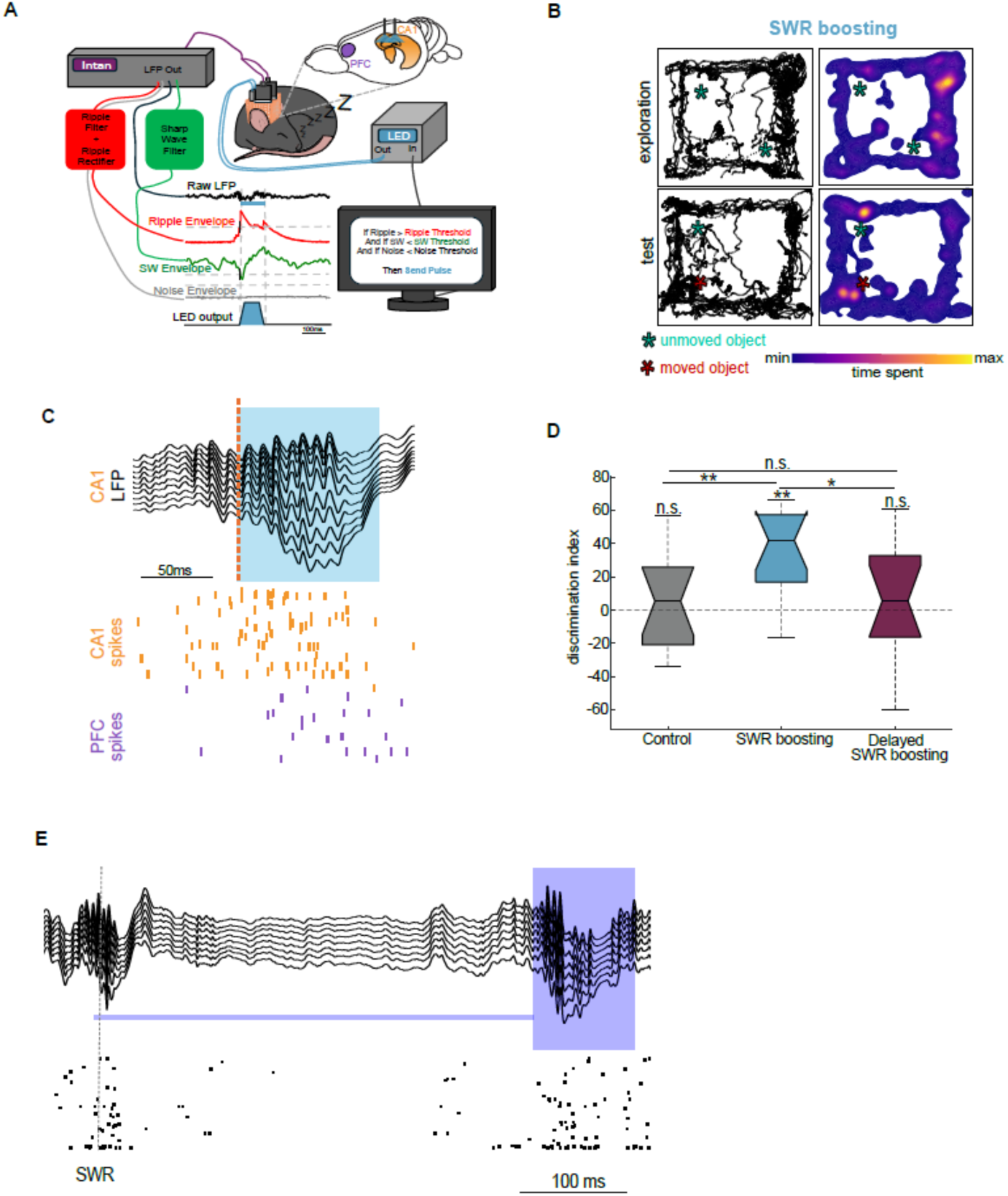
Closed-Loop setup and optogenetic manipulation. Related to Figure 3. **A**) Cartoon diagram of closed-loop optogenetic stimulation system. LFPs from CA1 pyramidal layer (“ripple channel”, red), CA1 stratum radiatum (Sharp-wave (SW) channel, green), and cortical channel (“noise channel”, grey) recorded through Intan. Ripple and Noise channels are band-pass filtered (100-300 Hz), rectified and low-pass enveloped calculated with a custom analog circuit^21^. SW channel is also band-pass filtered (10-40 Hz). A positive SWR detection is triggered when ripple and SW channels cross their respective pre-set thresholds, and the noise channel is below its threshold during the same time. A positive detection triggers the delivery of a trapezoidal current pulse to drive blue-light emission from two laser diodes. **B**) Top: Animal movement (left) and occupancy (right, heatmap) for a representative 5-minute training session (top) followed by SWR boosting during sleep and subsequent test (bottom). Cyan asterisks show the center of objects in the maze. Red asterisk represents a moved object during the test session. **C**) Example of optogenetic SWR boosting. Black trace is CA1 LFP, vertical dashed bar shows SWR real-time detection and blue rectangle shows optogenetic stimulation pulse. Raster plot below shows CA1 (in gold) and PFC (in purple) units firing. **D**) Memory recall performance (object discrimination index) for 5-minute training with no stimulation (same as Figure S3C), SWR-boosting (same as Figure 3F), and delayed stimulation sessions (control (grey): n = 13, p = 0.68, SWR boosting (blue): n = 13, p = 0.0017, delayed stimulation (magenta): n = 13, p = 0.74, Wilcoxon signed rank, control-SWR boosting: p = 0.0089, SWR boosting-delayed stimulation, p = 0.031, Wilcoxon rank sum test, control-delayed stimulation, p = 0.84, Wilcoxon rank sum test). **E**) Example LFP traces of a detected SWR (grey dotted line) followed by optogenetic stimulation (blue square) after a random delay (blue line). Below is raster plot of CA1 spikes (black tick mark, scale bar = 100ms).

**Figure S9.**
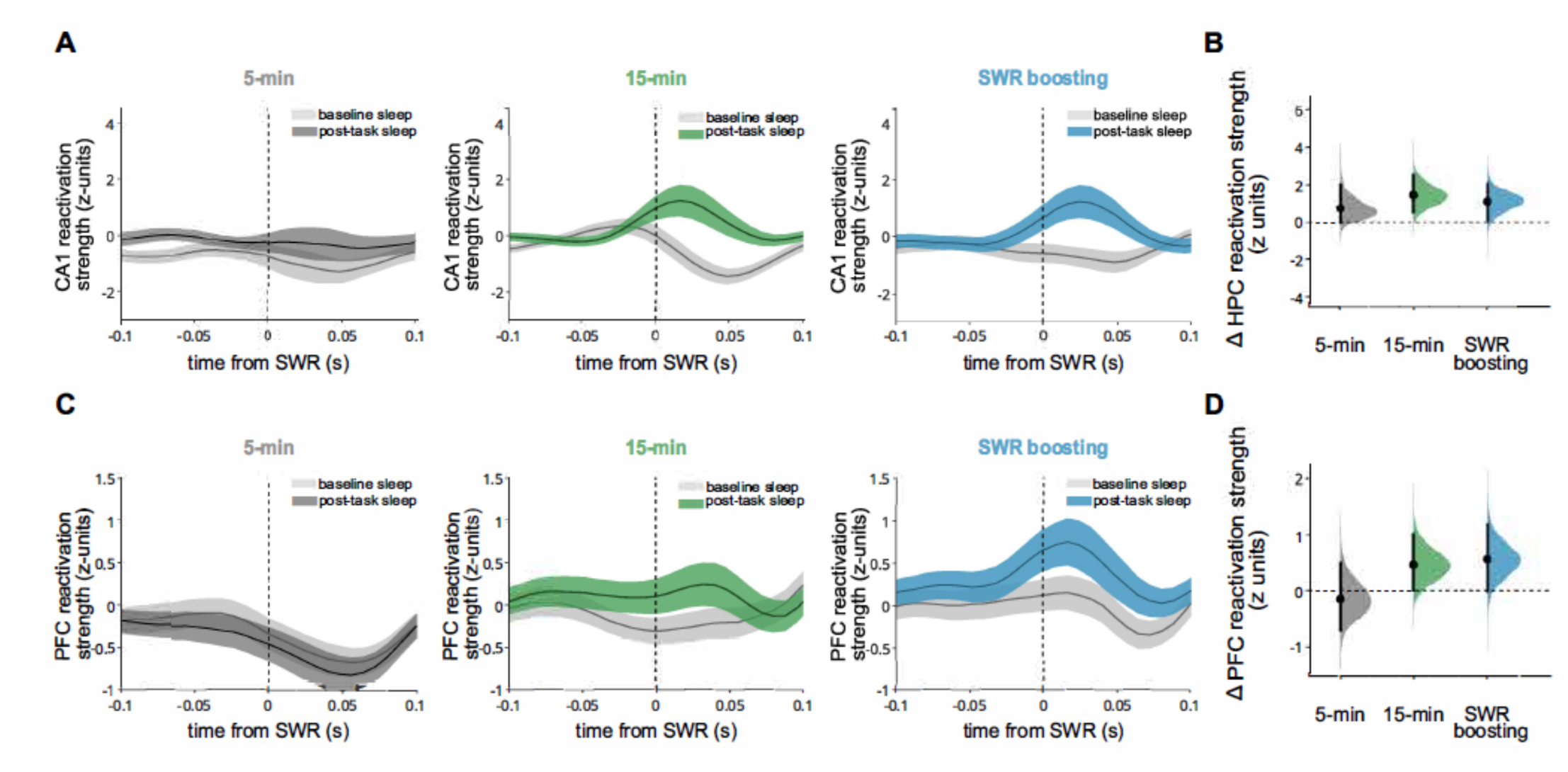
Reactivation analyses controlling for spiking increases in post-task sleep.Related to Figure 3 **A)** Same as Figure 3 but in this case spiking around SWRs was downsampled so mean firing rates in baseline and post-task sleep were matched (see Methods). **B)** Quantification of A) measured by hierarchical bootstrap estimate of assembly reactivation in post-task sleep relative to baseline sleep: 5-minute exploration (grey: P = 0.053, n = 97 assemblies); 15-minute (green: P = 6e-4, n = 189); SWR boosting (blue: P = 0.022, n = 131 assemblies). **C-D)** Same as A-B) but for PFC assemblies. 5-minute (grey: P = 0.68, n = 79 assemblies); 15-minute (green: P = 0.030, n = 96), SWR-boosting (blue: P = 0.032, n = 115).

**Figure S10:**
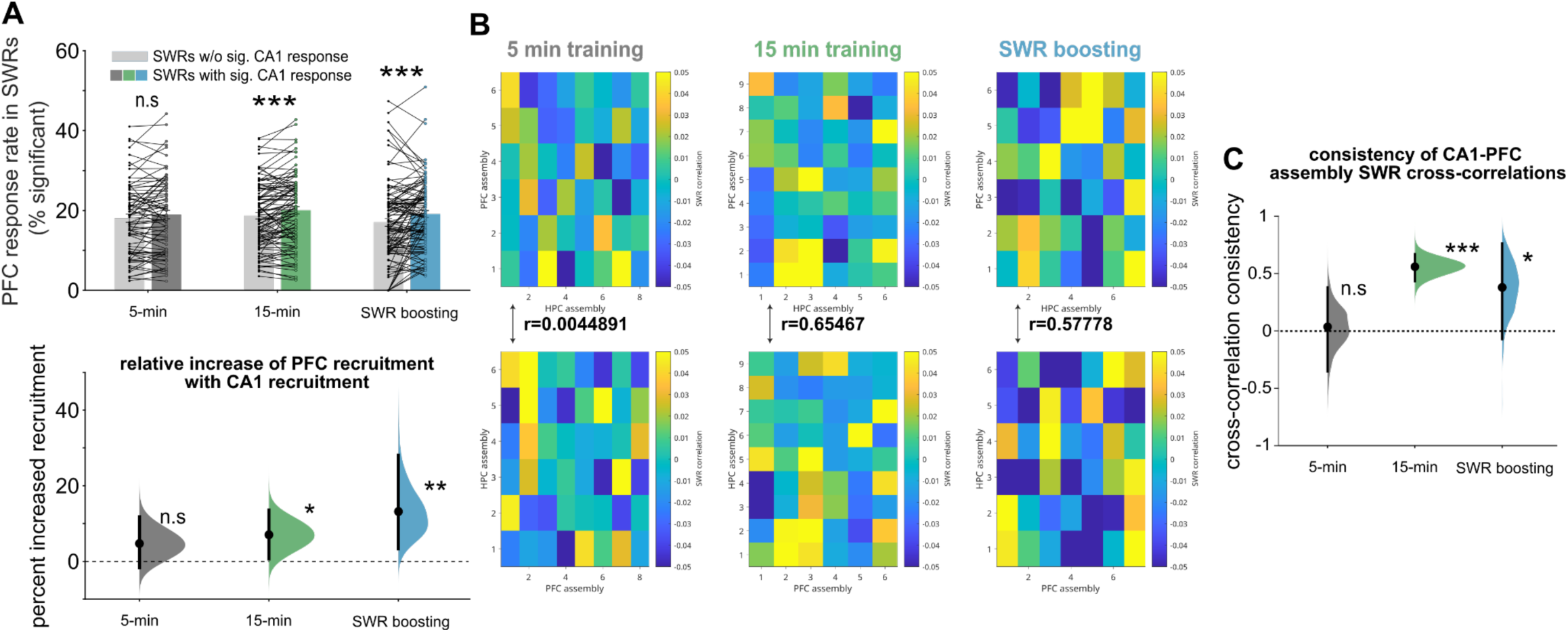
Coordinated reactivation of hippocampal and prefrontal assemblies. Related to Figure 4. **A)** PFC reactivation increase in SWRs with CA1 reactivation. Top: reactivation of prefrontal cell assemblies was similar following SWRs with (color) and without (light grey) significant CA1 assembly reactivation in sleep following 5-minute exploration (grey, P = 0.21, n = 85 assemblies, Wilcoxon signed-rank test), while following 15-minute exploration, PFC assemblies increased their reactivation in SWRs with significant CA1 reactivation (P = 8.7e-4, n = 103 assemblies, Wilcoxon signed-rank test), and SWR boosting reproduce this effect (P = 0.0041, n = 110 assemblies, Wilcoxon signed-rank test). Bottom: hierarchically bootstrapped estimate of the percent increase in PFC reactivation probability that is explained by CA1 reactivation; grey: 5 min training (P = 0.097, hierarchical bootstrap test); green: 15-minute exploration (P = 0.021, hierarchical bootstrap test); blue: SWR boosting (P = 0.0026, hierarchical bootstrap test). **B)** Example cross-correlation matrices between the reactivation in CA1 cell assemblies and the reactivation in PFC cell assemblies in SWRs in post-task sleep; SWRs have been split randomly into two halves (top: half1, bottom: half2). Note that the cross-correlation matrices remained highly consistent across the two halves of the data for 15-minute exploration sessions (Pearson’s r = 0.66, P < .00001, n = 54 assembly pairs) and 5-minute exploration with SWR boosting sessions (Pearson’s r = 0.58, P < 6.2e-5; n = 42 assembly pairs), but comparatively less for the 5 min training session (Pearson’s r = 4.5e-3, P = 0.98, n = 48 assembly pairs). **C)** Quantification of B) measured by hierarchical bootstrap estimate of the consistency of CA1 and PFC assembly reactivation in SWRs across post-task sleep: 5-minute exploration (grey: P = 0.43, hierarchical bootstrap test); 15-minutes (green: P = 0, hierarchical bootstrap test); SWR boosting (blue: P = 0.042, hierarchical bootstrap test).

## Notes

### Competing Interest Statement

The authors have declared no competing interest.

### Summary of Updates

New figures and text added to main manuscript and supplemental materials

